# Inferring population-level physiologically based model parameters for sleep in infancy and young childhood

**DOI:** 10.64898/2026.02.12.705640

**Authors:** Lachlan Webb, Andrew JK Phillips, James A Roberts

**Affiliations:** Brain Modelling Group, QIMR Berghofer, Herston, Brisbane, QLD 4006, Australia; Faculty of Health, Medicine and Behavioural Sciences, University of Queensland, QLD, Australia; Flinders Health and Medical Research Institute (Sleep Health), Flinders University, Bedford Park, South Australia, Australia

## Abstract

The sleep patterns of infants and young children differ from adult sleep patterns, with longer duration and multiple bouts per 24 hours. There is also considerable heterogeneity in infant sleep, both between individuals and within individuals across development. While mathematical models have been used to understand the mechanisms that regulate adult sleep, the development of sleep from infancy through early childhood has remained largely unexplored. Here we used an established mathematical model for adult sleep regulation to investigate how the underlying mechanisms of sleep mature from ages 1 month to 5 years, and identify a basis for inter-individual differences at each age. Using a Bayesian approach to estimate joint distributions of model parameters at different ages, we found that: (i) decreases in the rate of accumulation of sleep homeostatic pressure captured the reduction in sleep duration across development; while (ii) increases in the time scale of the sleep homeostatic drive captured the consolidation of sleep into fewer sleep bouts per day across development. In terms of inter-individual differences, we found larger spread in the parameters within the earlier stages of infancy (<1 year). The center and boundaries of the joint distributions evolved with age through the parameter space, converging towards previously described adult parameter values. A bifurcation analysis of the homeostatic time scale parameter revealed that progressive consolidation of sleep occurs through abrupt loss of bouts one at a time, punctuated by narrow intervals of non-entrained (>24-h) cycles. These results establish plausible, population-level trajectories in physiological parameters underpinning maturation of sleep regulation through infancy and early childhood.

**Author Summary:** The sleep patterns of infants and young children differ from adult sleep patterns, having both longer duration and being split into multiple bouts per 24 hours. Sleep patterns also vary greatly child to child and change considerably with development. While mathematical models have been used to understand adult sleep, the development of sleep from infancy through early childhood has remained largely unexplored. Here we used an established mathematical model for adult sleep regulation to investigate how the underlying mechanisms of sleep mature from ages 1 month to 5 years, and identify a basis for inter-individual differences at each age. Using our method, which inferred population level parameter distributions from population-level data summaries, we found that: (i) decreases in the rate of accumulation of sleep homeostatic pressure captured the reduction in sleep duration across development; while (ii) increases in the time scale of the sleep homeostatic drive captured the consolidation of sleep into fewer sleep bouts per day across development. In terms of inter-individual differences, we found larger spread in the parameters within the earlier stages of infancy (<1 year). These results establish plausible, population-level trajectories in physiological parameters underpinning maturation of sleep regulation through infancy and early childhood.

## Introduction

Infants and young children spend much more time asleep than adults. Sleep behaviors also change rapidly during these early years of development. Indeed, sleep patterns can be used as a marker of maturation and a milestone for development (Jenni & Carskadon 2012, Kocevska et al 2018). Emergence of circadian rhythms, night time sleep consolidation, and discontinuation of daytime sleep are all linked to increasing age (Bathory & Tomopoulos 2017, Jiang 2019, Shellhaas et al 2017). Longitudinal and cohort studies have quantified the maturation of population-level sleep characteristics (Iglowstein et al 2003, Jenni & Carskadon 2012, Mindell et al 2016, Staton et al 2020), showing that the amount of sleep per 24 hours and the number of daytime naps both decrease with age, while the lengths of sleep bouts increase. However, there is significant inter-individual variation in sleep maturation, with peak heterogeneity in sleep patterns during infancy (Jenni & Carskadon 2012). Deviations from typical-for-age sleep have been associated with later cognitive problems (Touchette et al 2007) and general health issues (Taveras et al 2008), and sleep disturbances are frequently comorbid with psychiatric conditions (Ivanenko & Johnson 2008). Sleep is thus deeply intertwined with human development and health. However, the underlying mechanisms in the subcortical sleep-wake circuitry that may contribute to the diverse sleep behaviors across development remain inadequately understood (Blumberg et al 2014).

Mathematical models of sleep that are based on physiology have been used to understand the underlying mechanisms that regulate sleep patterns in adults (Booth & Diniz Behn 2014, Fulcher et al 2014, Phillips & Robinson 2007, Postnova 2019, Tamakawa et al 2006) and nap consolidation in children (Athanasouli et al 2024). The Phillips and Robinson (2007) model has been extended to model sleep across a range of ages, including adolescence and older adults (Robinson et al 2011, Skeldon et al 2016, Skeldon et al 2017) and infancy (Webb et al 2024). There has long been a call for mathematical modeling of sleep across the lifespan (Crowley 2016) to improve understanding of both healthy sleep-wake behavior and sleep disorders in childhood (Jenni & LeBourgeois 2006). Moreover, most modeling attention has focused on identifying mean or typical parameter values, though this has received some attention in adults (Phillips et al 2010b, Skeldon et al 2021, Skeldon et al 2023). Previous work in infant sleep (Webb et al 2024) estimated individual developmental trajectories for model parameters for infants, with parameter trajectories that may not generalize to all children. It remains to be understood to what degree individual differences in child and infant sleep can be explained and generated within a single model. Understanding typical population ranges of parameters would provide new measures of inter-individual variation and additionally enable assessment of the extent to which abnormal data diverge from healthy.

Here, we aimed to elucidate the physiological underpinnings of infant and young child sleep, and identify how changes in those mechanisms are linked with development. To do this, we modeled the ascending arousal system across a range of post-natal ages from 1 month to 5 years. In particular, we used a Bayesian approach to identify age-specific distributions of biophysical parameters of the model (Phillips et al 2010a, Skeldon et al 2016) that explain the diversity in sleep behavior of children of a given age, and how these parameters evolve across development.

## Results

### Sleep-Wake Switch Model

We used a modified version of the sleep-wake model first presented in Phillips and Robinson (2007) (Figure 1). The 2007 version of the model is a neural mass model governed by three ordinary differential equations (ODEs) describing the monoaminergic (MA) and ventrolateral preoptic (VLPO) neuron populations and the sleep homeostatic drive. The model was later extended to include an additional three differential equations describing the coupling between an external light source and the natural circadian oscillator (see Forger et al (1999), Phillips et al (2010b) for more details). See Methods for more details.

**Figure 1:**
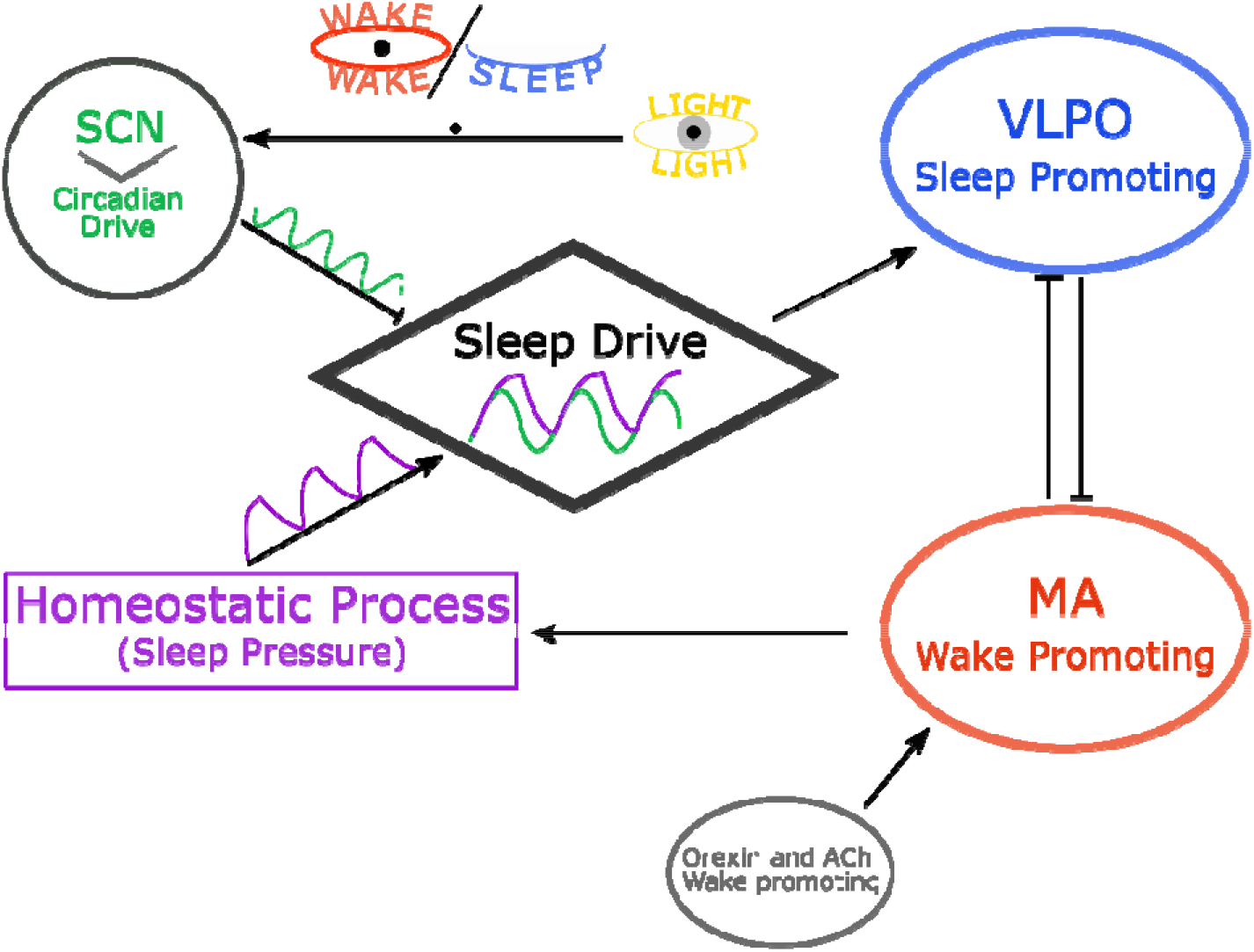
Schematic of the sleep model. The model describes the activity of the VLPO (ventrolateral preoptic nucleus, blue) and MA group (monoaminergic group, red), with mutual inhibition between them. The sleep-promoting VLPO receives input from the sleep drive, itself composed of a homeostatic drive (purple), a circadian drive (suprachiasmatic nucleus, SCN, green) entrained to light (yellow), where the light is gated by the activity state (wake versus sleep). The MA group receives input from Orexin and acetylcholinergic (ACh) neuron groups. Arrowheads denote excitatory influences, flat line ends denote inhibitory influences.

This model has been well validated (Phillips et al 2010a, Phillips & Robinson 2008, Postnova et al 2012, Puckeridge et al 2011, Skeldon et al 2017) and has been used to understand changes in sleep across the human lifespan (Skeldon et al 2016, Webb et al 2024). Thus, it is a natural starting point for understanding the diversity and maturation of sleep in infancy and early childhood.

### Population Estimates of Sleep Behavior

We sought to explain two key features of infant and young child sleep: the long duration of sleep, and its polyphasic nature (day time naps and night wakings). Hence, we fitted two key sleep behavior characteristics: the average amount of sleep per day and the average number of sleep bouts per day. We did this at the population level (meaning all children of a certain age), aiming to fit distributions rather than point estimates. We used reported summary statistics from meta-analyses and cohort studies (Dias et al 2018, Iglowstein et al 2003, Jenni & Carskadon 2012, Sadeh et al 2009, Staton et al 2020) (Appendix S1) to construct probability distributions at seven different ages: 1, 3, 6, and 9 months old, and 1, 3, and 5 years old (Figure 2; see Methods). When constructing distributions for average number of bouts of sleep per day, we defined a bout of sleep as including: i) a single, consolidated nighttime sleep, ii) a daytime nap, or iii) a period of sleep between nightwakings/daytime wake (e.g., a 24-h period that had one daytime nap and one night waking within the night sleep would be characterized as having three bouts of sleep). Bouts in the sleep model output were defined as any periods of continuous sleep.

**Figure 2:**
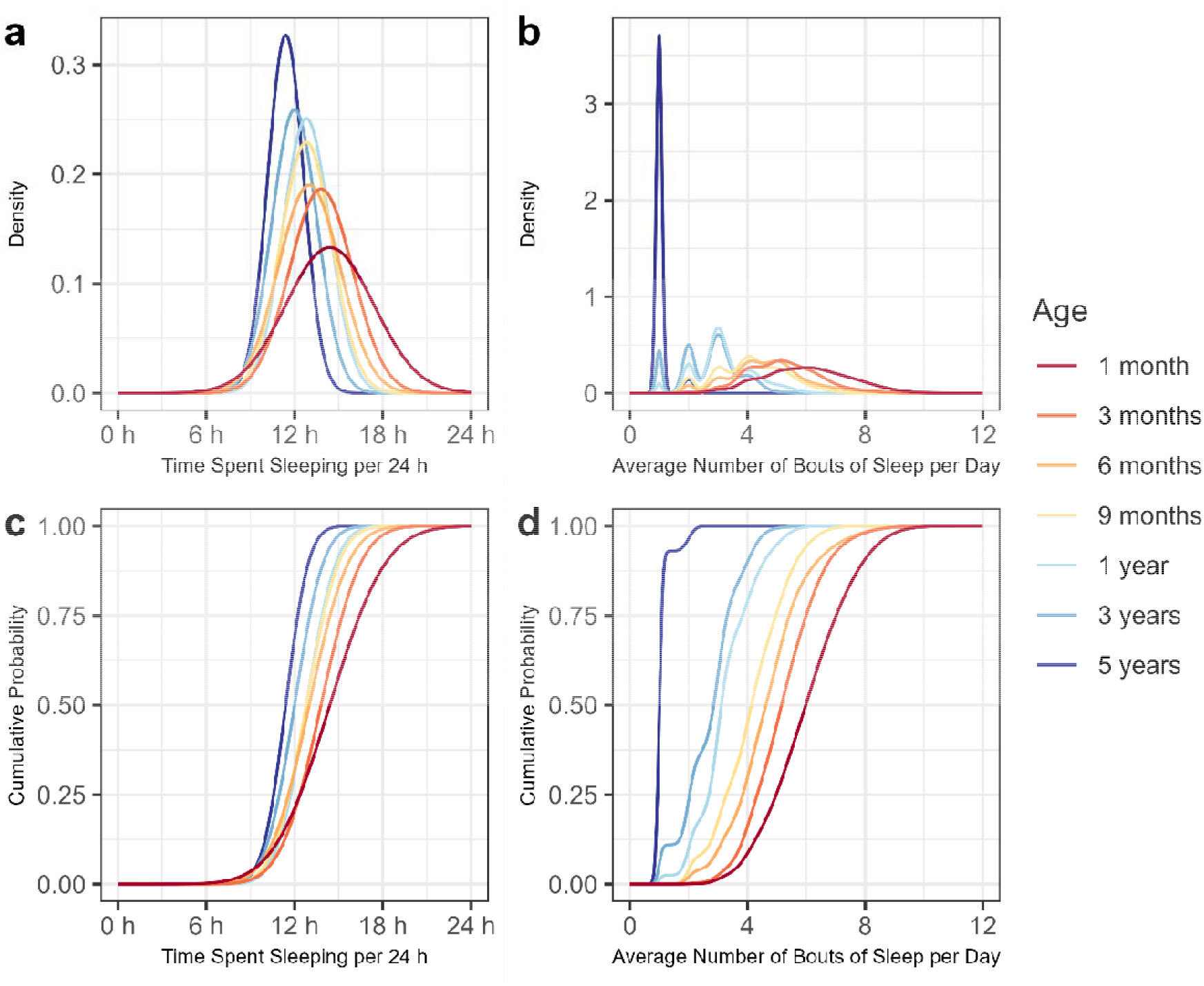
Empirical distributions of daily sleep duration and bout numbers as a function of age. (**a**) The density for amount of sleep per 24 hours for each age. (**b**) The density for average number of bouts of sleep per 24 hours for each age. (**c**) The cumulative density of sleep per 24 hours for each age. (**d**) The cumulative density for average number of bouts of sleep per 24 hours for each age. The empirical population for the average amount of sleep per 24 hours, (a) and (c), is assumed normally distributed (Table 2). A mixed Gaussian distribution is used for the average number of bouts per 24 hours, (b) and (d), and the parameters for the mixed Gaussian distributions are chosen such that the distributions are consistent with the literature; e.g., proportion not napping, proportion having 1 nap per day, etc. (Table 2 and Appendix S1).

**Table 1:** Parameter values used as a baseline at maturity.

| Sleep/wake regulation |  |  |  | Circadian |  |  |  | Light |  |
| --- | --- | --- | --- | --- | --- | --- | --- | --- | --- |
| Parameter | Value | Parameter | Value | Parameter | Value | Parameter | Value | Parameter | Value |
| $Q_{\max}$ | 100 s<sup>-1</sup> | $v_{vc}$ | -2.9 mV | $\alpha_0$ | 0.16 | $\kappa$ | 12/$\pi$ h | $c$ | 1/6000 |
| $\theta$ | 10 mV | $v_{vh}$ | 1.0 mV nM<sup>-1</sup> | $I_0$ | 9500 lux | $\gamma$ | 0.23 | $l_e$ | 0 lux |
| $\sigma'$ | 3 mV | $D_0$ | -10.2 mV | $p$ | 0.6 | $f$ | 0.99669 | $l_d$ | 500 lux |
| $A$ | 1.3 mV | $\chi$ | 45 h | $b$ | 1.0 | $\tau_c$ | 24.2 h | $s_1$ | 8 h |
| $v_{vm}$ | -2.1 mV s | $\mu_{hm}$ | 4.4 nM s | $G$ | 19.9 | $k$ | 0.55 | $s_2$ | 17 h |
| $v_{mv}$ | -1.8 mV s | | | $\lambda$ | 60 | | | | |
| $\tau_{m,j}$ | 10 s | | | $\beta$ | 0.013 | | | | |

**Table 2:** Distribution parameter values used for the probability model for sleep characteristics versus age.

| Age | Sleep |  | Bout |  |  |
| --- | --- | --- | --- | --- | --- |
| | Average ($\mu$) | Standard Deviation ($\sigma$) | Average ($\mu$) | Standard Deviation ($\sigma$) | $\Phi$ |
| 1m | 14.4 h = 0.600 day | 3.00 h = 0.125 day | 6.005 | 1.4542 | [0,0.001,0.025,0.111,0.021,0.29,0.22,0.133,0.01] |
| 3m | 13.8 h = 0.575 day | 2.14 h ≈ 0.089 day | 5.207 | 1.2154 | [0,0.004,0.04,0.22,0.34,0.3,0.07,0.025,0.001] |
| 6m | 13.0 h ≈ 0.542 day | 2.10 h ≈ 0.088 day | 4.675 | 1.324 | [0,0.04,0.11,0.28,0.375,0.12,0.05,0.025,0] |
| 9m | 12.8 h ≈ 0.533 day | 1.74 h ≈ 0.073 day | 4.135 | 1.166 | [0,0.08,0.19,0.345,0.285,0.1,0,...] |
| 12m | 12.8 h ≈ 0.532 day | 1.59 h ≈ 0.066 day | 3.225 | 0.968 | [0.025,0.15,0.5,0.225,0.1,0,...] |
| 36m | 12.0 h = 0.500 day | 1.54 h ≈ 0.064 day | 2.73 | 0.959 | [0.11,0.25,0.45,0.18,0.1,0,...] |
| 60m | 11.4 h ≈ 0.475 day | 1.22 h ≈ 0.051 day | 1.07 | 0.2778 | [0.93,0.07,0,...] |

Motivated by previous studies, we focused on two-parameter fits to these age-dependent distributions. Previous works have used the rate of increase in homeostatic drive from the MA group, *μ*_ℎ*m*_, to explain the changes in amount of sleep across infancy (Webb et al 2024) and adolescence (Skeldon et al 2016). Multiple bouts of sleep per day have previously been explained by varying the time scale of the homeostatic drive, *χ*, in both infants (Webb et al 2024) and other mammals (Phillips et al 2010a). We therefore estimated age-dependent parameters ***ψ_i_*** where ***ψ_i_*** for each age *i* is a vector [*μ*_ℎ*m*_, *χ*]. Other works have considered a different parameter, the constant inhibition to the VLPO *D*_0_ (Phillips et al 2010a), instead of *μ*_ℎ*m*_ to explain the varying amount of sleep, which we consider in the Supplementary Material (Appendix S2).

### Bayesian Model Results

Results of the parameter fitting showed that at the younger ages (1-9 months old), *μ*_ℎ*m*_ exhibited wide distributions with higher central values, and lower centered, narrower distributions at the older ages (Figure 3a). The posterior distributions for *log χ* shifted higher with age (Figure 3b), with the distributions for the monophasic sleeps in the 1+ year old ages hitting the assumed upper boundary (adult value) of *ln χ*.

**Figure 3:**
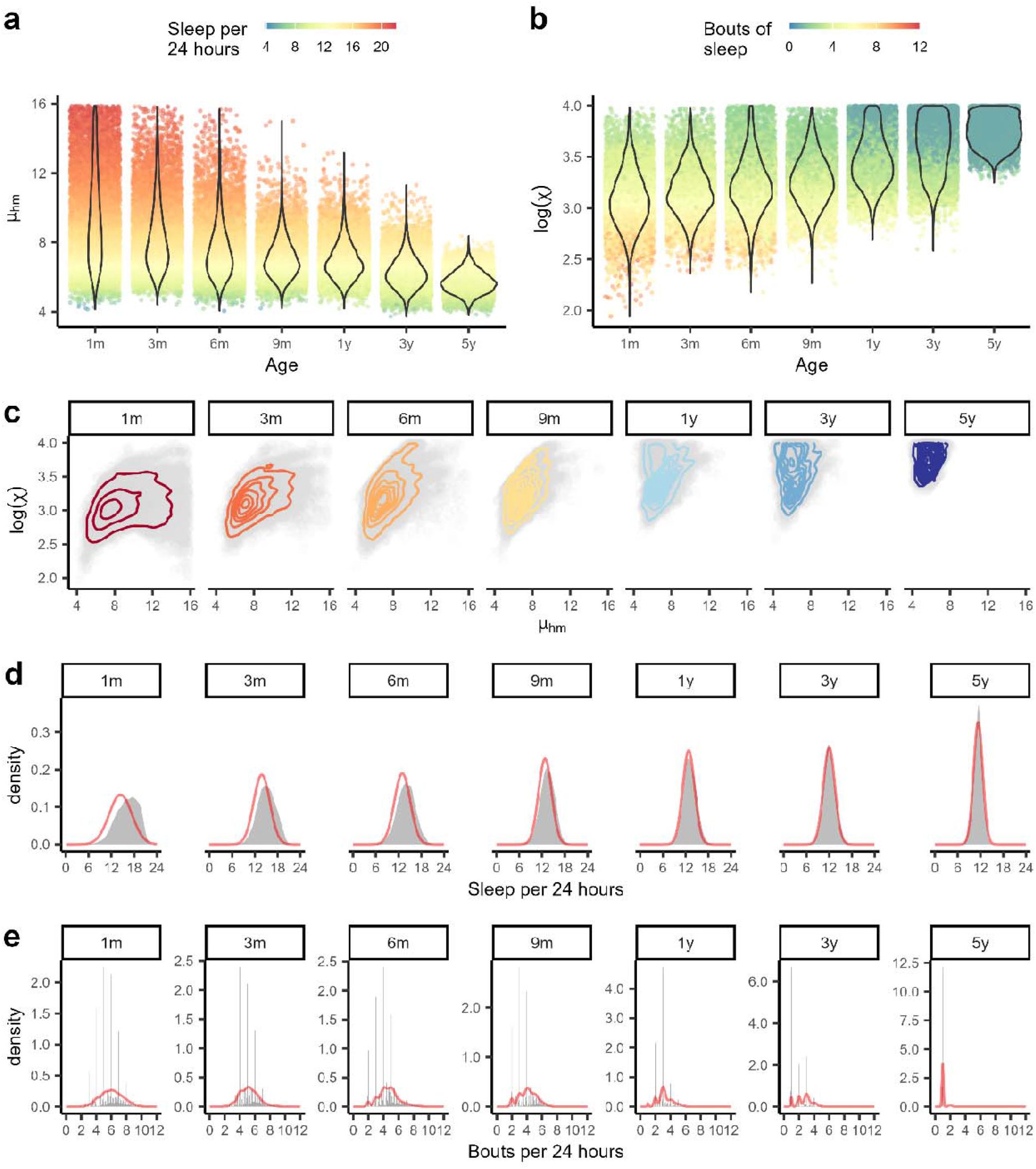
Age-specific parameter estimation in the *μ*_ℎ*m*_ − *χ* model. **a)** Posterior distributions (violins) for each age for *μ*_ℎ*m*_, with points colored by the corresponding average sleep per 24 hours. **b)** Posterior distributions (violins) for each age for log *χ*, with points colored by the corresponding number of bouts of sleep. **c)** The joint posterior distribution (gray) for each age with density contours shown at density levels 0.05,0.01,0.15, **d)** Distributions by age for the amount of sleep per 24 hours arising from the parameter posteriors (gray) superimposed with the empirical density curves (red). **e)** Distributions for the average number of bouts per 24 hours.

There was a near one-to-one relationship between *μ*_ℎ*m*_ and total sleep per day, as seen from the smooth color gradient (Figure 3a). In contrast, the relationship between bouts and *log χ* is modulated by *μ*_ℎ*m*_, as seen by the blend of colored points in Figure 3b, and the diagonal boundary in Figure 3c between the polyphasic and monophasic area in the parameter space (in particular the boundary between the almost completely monophasic 5-year-old contour and the almost completely polyphasic 9-month-old contour). This means the values of *χ* needed to set the average number of bouts per 24 hours will depend on the value of *μ*_ℎ*m*_, and consequently the amount of sleep per 24 hours.

At the older ages when sleep becomes monophasic, the posterior distributions of *log χ* hit the upper *log χ* boundary. This is due to the relationship between *χ* and number of bouts having a large range of values where only one bout is produced. Once *χ* is in that range, the number of bouts becomes an ineffective constraint on the dynamics. Nevertheless, although the joint posterior extends into those higher *χ* spaces (Figure 3c), contours of the distributions show that the bulk of the weight lies away from the upper bound on *χ*.

The model shows good agreement comparing empirical to fitted sleep characteristics (Figures 3d, e), though younger ages exhibit a slight skew towards a high amount of sleep per 24 hours (Figure 3d). The model sleeps have a more discrete distribution of number of bouts than assumed in the population, and the model overestimates the probability of having a single bout of sleep per day in the older age groups (Figure 3e).

Plotting the joint parameter distributions, ***ψ_i_***, on the same axes reveals the age trend more directly (Figure 4). The center of density curve exhibits a well-behaved trajectory through the parameter space across development. The center and edges shift upwards (to higher *log χ*) and to the left (to lower *μ*) with increasing age. There is little difference in the *μ*_ℎ*m*_ range of the contour between 9 months and 1 year of age, which is expected as the population data had the same mean amount of sleep per 24 hours for both ages.

**Figure 4:**
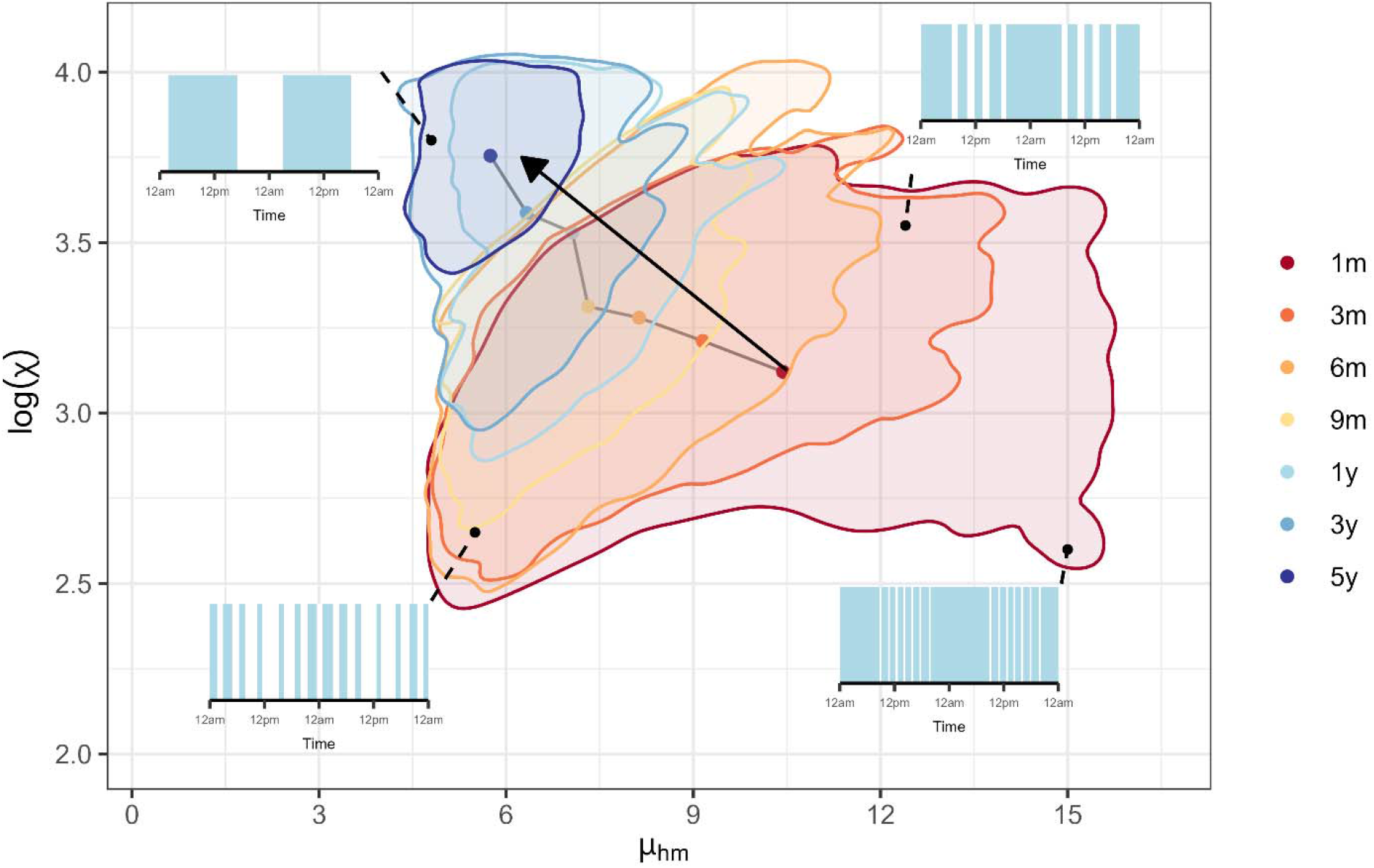
Contours from a kernel density estimate containing 95% of the joint posterior sampled points of *μ*_ℎ*m*_ and log *χ* posteriors at each age. Centers of distributions are shown with dots, colored by age; arrow denotes the direction of increasing age. Insets show examples of modelled sleep behavior at the parameters indicated by black dots, with light blue bars signifying bouts of sleep.

Significant overlap of the contours is consistent with the significant heterogeneity in sleep characteristics observed across the population. Examples of sleep behavior (Figure 4 insets) illustrate the diversity of patterns consistent with a single age and/or across ages.

### Bifurcations in sleep patterns

We next sought to understand the dynamical origins of the relationship between parameters and sleep characteristics, and more broadly how the inferred age-related changes in parameters change the model dynamics. We examined time series of the model as a function of *χ* and *μ*_ℎ_, varying one at a time and extracting sleep/wake patterns, total sleep durations per day, numbers of bouts per day, and local extrema of the homeostatic drive *H*. Decreasing *χ* results in abrupt transitions from consolidated sleep to increasingly many bouts of sleep (Figure 5). Bouts of sleep disappear/emerge at clock times distinct from existing bouts (Figure 5a), in concordance with the expected behavior of napping children—namely that a child moving from (say) two naps to one nap does not have the naps coalesce but rather drops one or the other. For this parameter set, almost all bouts of sleep occur between 2pm and 7am (i.e., primarily the night). With increasing *χ*, the bout that is lost at each transition is the final bout before the long day-time wake period. Most of the range of *log χ* has an integer number of bouts per 24 hours, but there are also narrow sections of non-integer numbers of bouts—occurring at transitions between integer regimes—where the sleep pattern does not repeat on a 24-hour period (Figure 5b and c). This implies increased diversity of napping patterns at the boundary between two napping regimes, which is consistent with common experience (Davis et al 2004).

**Figure 5:**
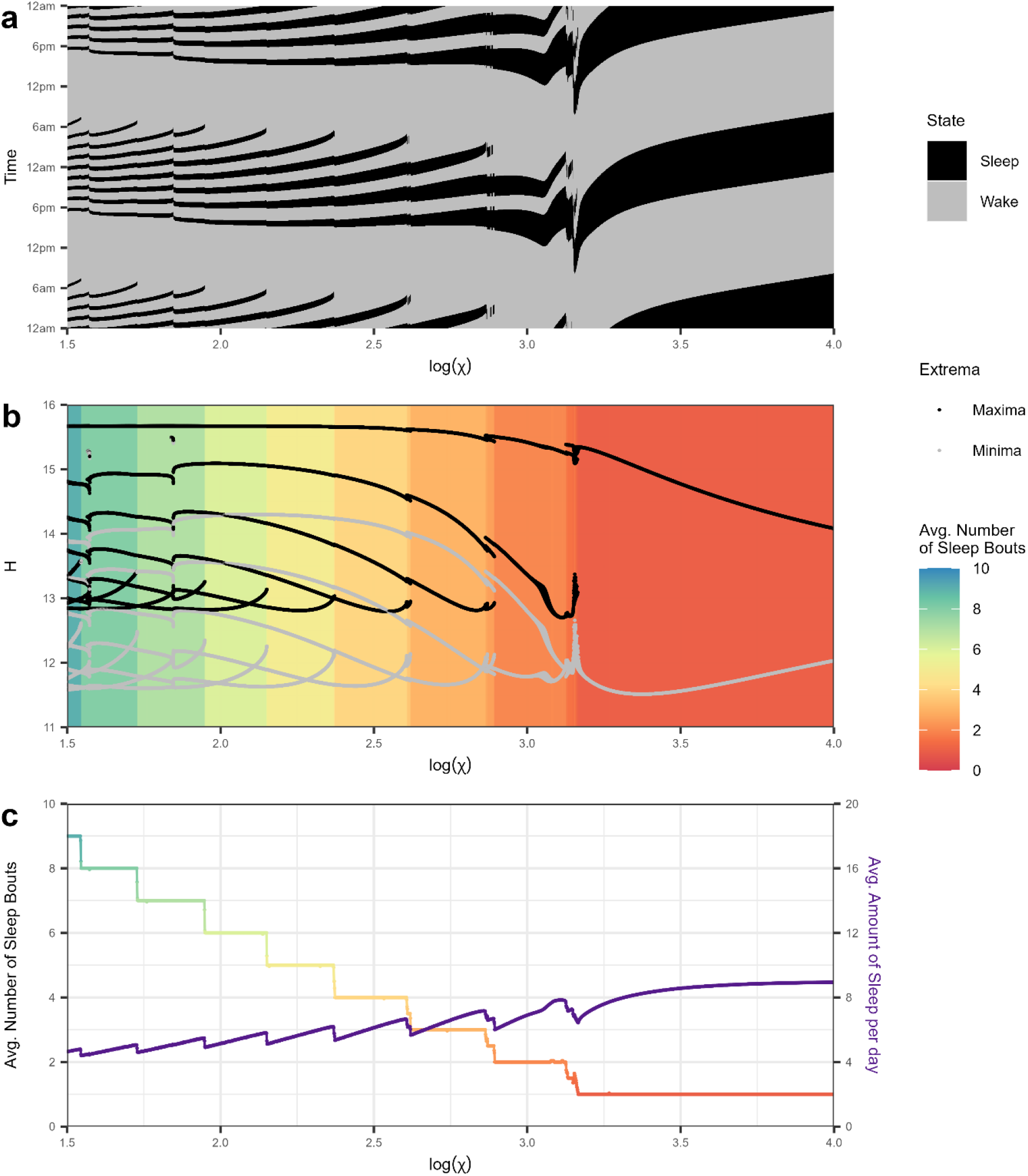
Dynamics of homeostatic drive and sleep behavior with changing sleep model parameter *χ*. **a)** The sleep-wake behavior across 2 days from different values of log *χ*, where black is sleep and gray is wake. **b)** Bifurcation plot of homeostatic drive and average number of sleep bouts across 7 days of sleep-wake activity as a function of log *χ*. Black dots denote local maxima of the homeostatic drive and gray dots the local minima. Colored shading shows the average number of bouts per 24 hours, highlighting values of *χ* that produce integer values of average number of bouts per day, interspaced with values that produce non-integer values. **c)** Average number of bouts of sleep per day (left axis, colored as b) and average amount of sleep per day (right axis, purple) across 7 days of sleep-wake activity as a function of log *χ*.

We also examined the changing dynamics of the sleep homeostatic drive with changes in the parameters of interest, as this gives insights into how the internal brain chemistry of the sleep-wake switch changes with likely trajectories of neurodevelopment. In the one-bout domain (*log χ* > 3.2), increasing *χ* results in a smaller amplitude of the homeostatic drive (Figure 5b) and minimal changes to the amount of sleep per day (Figure 5c). This implies that the range of the somnogen levels in the homeostatic drive decreases as the time scale (*χ*) approaches the mature, adult value.

Increasing *μ*_ℎ*m*_ also increases the number of bouts of sleep. Figure 6a shows the corresponding sleep state behavior from increasing *μ*_ℎ*m*_, with a higher *μ*_ℎ*m*_ causing wake to occur primarily between approximately 12am - 12pm, and wake time and sleep onset time shifting earlier before multiple sleep bouts per day appear. Changes in the dynamics of the homeostatic drive from increasing *μ*_ℎ*m*_ are shown in Figure 6b-c, with the amount of sleep per day and number of bouts per day increasing as *μ*_ℎ*m*_ rises. There are large ranges of *μ*_ℎ*m*_ where the average number of bouts of sleep per day is non-integer (Figure 6c), that also correspond with values of *μ*_ℎ*m*_ where small perturbations in the parameter can have dramatic impacts on the sleep patterns (Figure 6a). Decreasing *μ*_ℎ*m*_ in the 1 bout space also reduces the dynamical range of the homeostatic drive (Figure 6b), similarly to increasing *χ* in Figure 5b. However, in both converse cases (decreasing *χ* and increasing *μ*_ℎ*m*_) the dynamical range of the homeostatic drive remains roughly the same.

**Figure 6:**
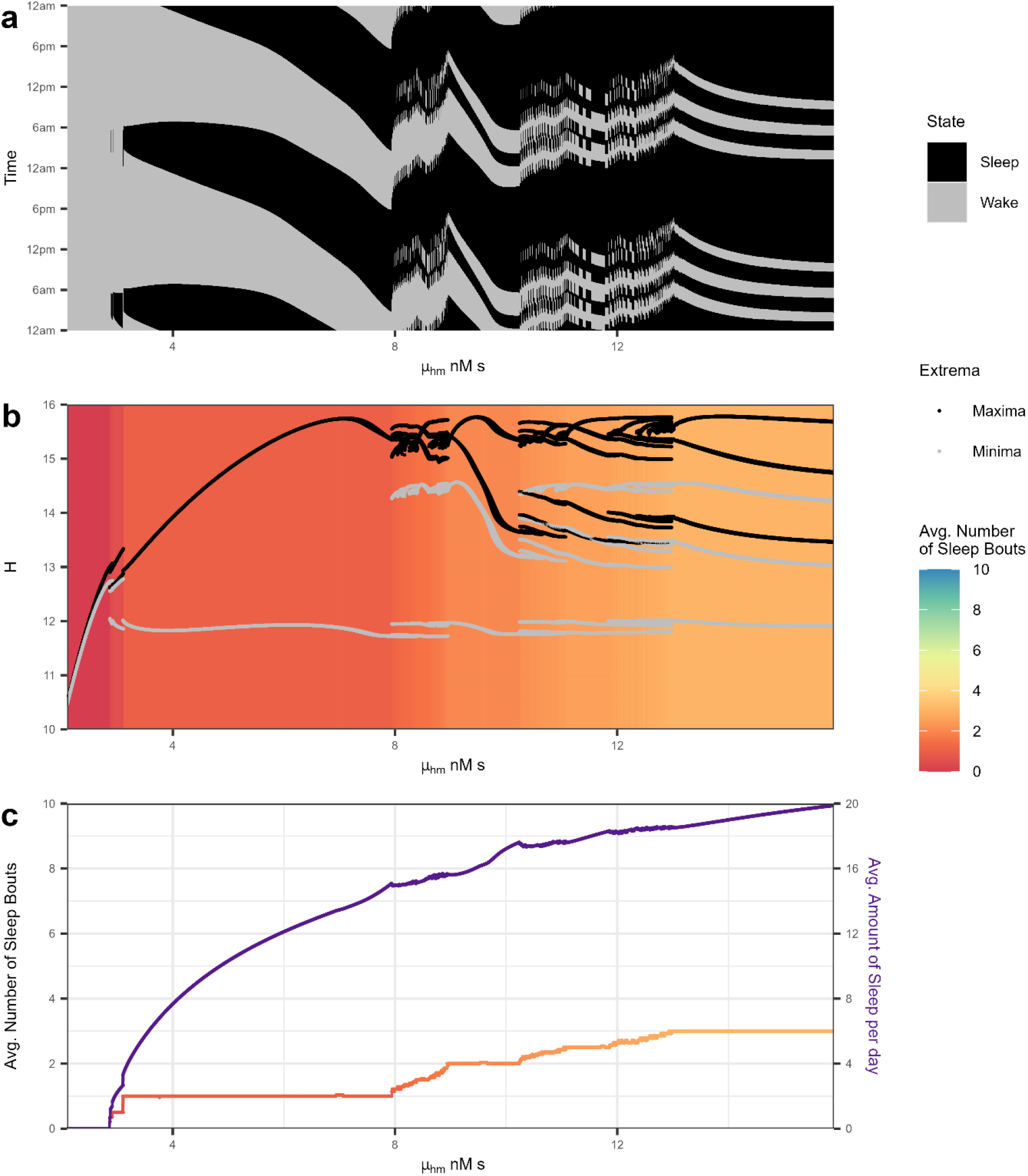
Dynamics of homeostatic drive and sleep behavior with changing sleep model parameter *μ*_ℎ*m*_. a) The sleep wake activity across 2 days for a range of *μ*_ℎ*m*_, where black is sleep and gray is wake. **b)** Bifurcation plots of homeostatic drive across 7 days of sleep-wake activity as a function of *μ*_ℎ*m*_ , showing local maxima (black) and local minima (gray) of the time series. Colored shading (same color scale as Figure 5b) shows the average number of bouts per 24 hours, showing how increasing *μ*_ℎ*m*_ produces multiple bouts of sleep per day. **c)** Average number of bouts of sleep per day (left axis, colored as b) and average amount of sleep per day (right axis, purple) across 7 days of sleep-wake activity as a function of *μ*_ℎ*m*_.

The relationship between the timing of the sleep bouts (Figure 5a and 6a) and homeostatic drive dynamics (Figure 5b and 6b) can be viewed directly in the time series of the homeostatic drive (Figure 7). With low *χ*, the nighttime sleep was broken into many bouts (Figure 5a) which corresponds to sawtooth-like activity on the decreasing side of the homeostatic drive waveform (Figure 7a and b). In contrast, higher *μ*_ℎ*m*_ introduced morning naps (after the main sleep, Figure 6a) with sawtooth-like activity in the increasing side of the homeostatic drive waveform (Figure 7c and d). When both parameters are in an infant space (c.f. Figure 4) the sawtooth like activity is throughout the homeostatic drive waveform (Figure 7e), as was theorized for the infant homeostatic drive in Jenni and LeBourgeois (2006).

**Figure 7:**
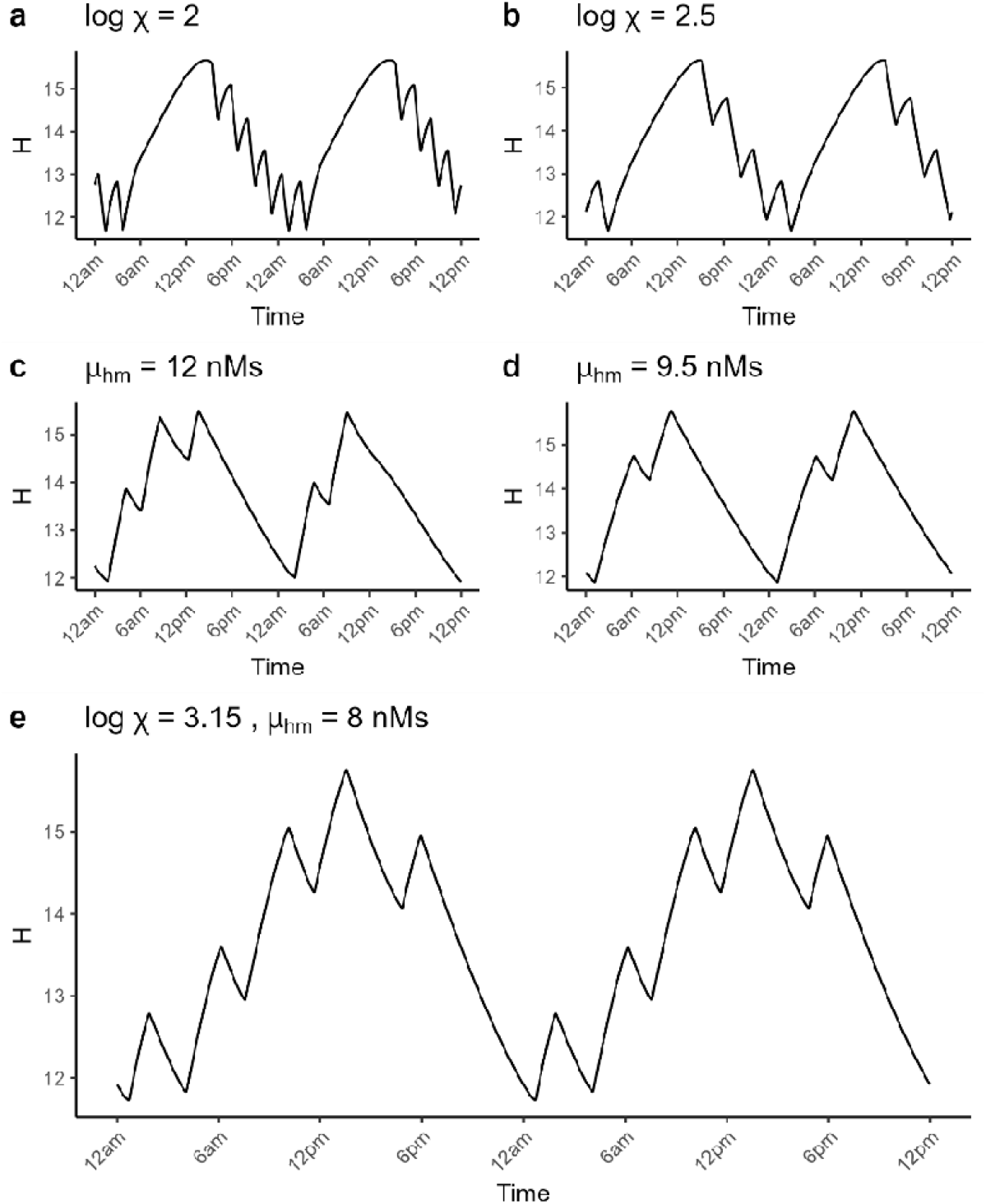
Time series of the homeostatic drive under different vales of *χ* and *μ*_ℎ*m*_. **a)** The homeostatic drive across 2 days when log *χ* = 2, and **b)** when log *χ* = 2.5. **c)** The homeostatic drive across 2 days when *μ*_ℎ*m*_ = 12 nMs, and **d)** when *μ*_ℎ*m*_ = 9.5 nMs. **e)** The homeostatic drive across 2 days when log *χ* = 3.15 and *μ*_ℎ*m*_ = 8 nMs. Unless specified, all other parameters are set to values given in Table 1.

### Further comparisons to real world populations

We next sought independent tests of the model against other summary measures from published data, to which the model was not explicitly fitted. Examining if the model fitted to a set of primary sleep characteristics (sleep amount and number of sleep bout lengths) consequently produces distributions consistent with independent data on secondary statistics makes this a strict test of our model. The Bayesian analysis produces a time series of sleep wake behavior for each set of sampled parameters ***ψ_i_***, which can be further summarised beyond sleep per day and bouts per day. For example, Mindell et al (2016) presented data across age (collected using a sleep diary app) on distributions of sleep bout lengths and the relationship between bout length and the clock time of sleep onset (Figure 8a-b). These data show (i) a transition from a skewed distribution to a clear bimodal distribution in bout lengths across the first year of life (Figure 8a), and (ii) a complex relationship with clock time with considerable diversity in sleep onset times and the higher-mode of long sleeps emerging as a distinct “island” in the plot (Figure 8b). We emulated this analysis on our model-generated sleep time series. Specifically, we summarized the length and timing of the complete sleeps from the last seven days of each 35-day time series, and compared these ‘synthetic populations’ of behavior (Figure 8c and d) to the published data of Mindell et al (2016).

**Figure 8:**
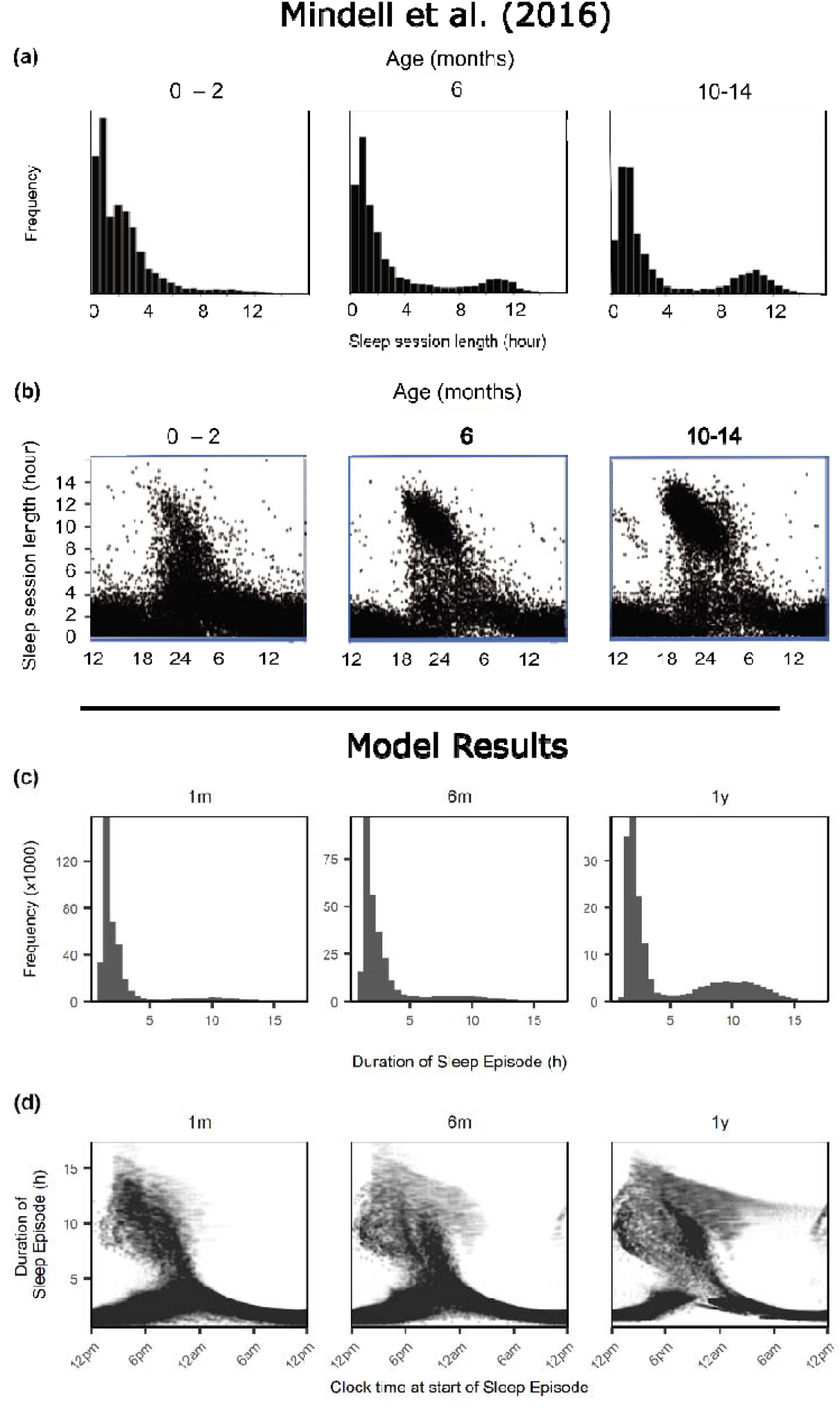
A comparison between Mindell et al (2016) sleep characteristics across ages and model results. (**a**) Empirical histograms of sleep session lengths for the age ranges 0-2 months, 6 months, and 10-14 months of age. (**b**) Scatter plots of sleep session length by the time of the start of the sleep bout for the same age ranges as panel a. Compared to modelled sleep bout duration distributions and sleep bout onset timing across development; the characteristics of the sleeps in the time series produced by model posterior draws for ages 1 month, 6 month and 1 year old. (**c**) Histograms of the length of each sleep bout from model posterior draws for ages 1 month, 6 month and 1 year old. (**d**) Scatter plots of sleep bout duration versus the clock time of the start of the bout for the same ages as c. Panels a and b adapted from Mindell et al (2016).

The histograms in Figure 8c show the shifting distribution of many short bouts of sleep (at 1 month of age) to a bimodal distribution of some long bouts and some short bouts, a trend also observed in the Mindell et al (2016) figure (Figure 8b), to more episodes of consolidated sleep lasting 8-12 hours for the 1-year age. The scatter plots of bout duration versus the clock time of bout start in Figure 8d for the younger ages show short sleeps commencing throughout the day, with the longer sleeps commencing in the afternoon and evening, again similar to the data. In the 1-year panel in Figure 8d, there is a lack of short (<2 h) bouts commencing in the night, showing the consolidation of night time sleep. Both the model and the data exhibit a wide range of sleep onset times, and exhibit very similar overall structure. The model long sleeps have a wider range of onset times. While the majority of sleeps have evening onset, a small proportion of the 10+ hour sleeps start at any time of the day, particularly for the 1-year panel; unsurprising given that we have not attempted to fit to sleep timing. This suggests that additional constraints on the circadian alignment may be necessary to capture sleep onset timing. This is confirmed in the Supplementary Material, where the displaced sleep episodes are shown to correspond to non-entrained dynamics (over 17% of 1-year-olds with a period >24.2h) indicating a lack of entrainment to the solar curve (Appendix S3). This lack of entrainment accumulates throughout a long simulation to produce ill-timed sleep bouts.

We further examined the effect of the parameter *b* (Appendix S4), the phase response of the circadian pacemaker to the solar curve. This parameter had the fixed value of 1 in all results, based on previous findings in modeling infant sleep (Webb et al 2024). Changing this parameter towards accepted adult values had negligible impact on the joint *χ* − *μ*_ℎ*m*_ distribution, though did shift the circadian dynamics closer to achieving 24 h entrainment (Supplementary Figure S7 and Supplementary Figure S8).

Nevertheless, even when not constrained to do so, the model reproduced the key features of sleep bout timing expected for young ages, primarily small sleeps spread throughout the day with some longer sleeps at night, and the shift to consolidated night sleep with a bimodal distribution of sleep bout lengths.

## Discussion

Sleep develops rapidly in the first few years of life, yet the physiological mechanisms underpinning these changes remain elusive. Here, we used a computational model to infer how physiological processes develop during the early years of human life. We used Bayesian parameter estimation to understand how the heterogeneous sleep characteristics of a population are linked to distributions of underlying physiological parameters, and how these distributions interact and evolve with age. Our findings reveal key changes in sleep homeostatic regulation that may underlie developmentally linked changes in both sleep duration and sleep consolidation from ages 1 to 5 years old, and the differences in that regulation underpinning the heterogeneity of sleep behavior at a given age.

We explored the parameter combination of *μ*_ℎ*m*_ (the increase in homoeostatic drive from the wake promoting MA group) and *χ* (the time scale of the homeostatic drive) to explain the heterogeneity and change in sleep behavior across infancy and early childhood. The resulting parameter posteriors exhibited a decrease with age in the spread and center of the distribution for the parameter largely controlling amount of sleep (*μ*_ℎ*m*_), while the parameter distribution for *χ* shifted upwards with age, coinciding with the increasing consolidation of sleep across development. These results broadly align with the parameter trajectories observed in the longitudinal, single-N approach in Webb et al (2024). Similar results were found when we used a different parameter to control the amount of sleep per day (Appendix S2).

Although our focus is on infant and child sleep, the model and Bayesian inference method are general and applicable to other ages and cohorts. We have demonstrated proof of principle that population summaries of sleep characteristics can be used to build probability models and find posterior distributions of underlying parameters of the sleep regulation network. For example, one could find summary sleep characteristics for neurological conditions that have sleep disturbance as a part of the pathology, estimate parameter posteriors, and compare these with parameters for typical sleep to identify the key physiological differences.

In the present work we have estimated joint posteriors for only two parameters at a time. Expanding the framework to produce posteriors for more than two parameters is nontrivial without additional orthogonal data features to constrain the fitting. For example, some of the Phillips-Robinson model parameters have complementary relationships with the sleep characteristics, such as *μ*_ℎ*m*_ and *D*_0_ both directly influencing the amount of sleep. We could only include one of these parameters in ***ψ_i_*** as changes in one parameter could be approximately counteracted by reverse changes in the other. Moreover, adding more parameters to the probability model increases the computational time.

To characterize sleep-wake patterns, we used two summary statistics: sleep duration and number of sleep bouts per day. Different measures may more accurately characterize sleep behavior at other ages. For example, Skeldon et al (2016) used sleep onset/offset and sleep midpoint when modeling the changes in sleep behavior from adolescence to old age, since sleep is generally monophasic in this age range and therefore sleep bouts per day is a relatively uninformative summary statistic. Adding more sleep behavior characteristics and allowing more sleep-wake model parameters to vary in the probability model increases the complexity and run time. Hence, we constrained our analysis to sleep duration and sleep bouts, as these exhibit the greatest variability in our modelled age range, and the parameters to *μ*_ℎ*m*_ and *χ* as they have been shown to closely relate to those sleep characteristics (Webb et al 2024). To use this Bayesian framework across the entire lifespan, it may be most effective to develop models with age-specific characteristics in the probability model.

We explored parameters in the sleep pressure drive, but did not deeply explore changing other parts of the model. For example, it has been suggested that the parameter *b* in the circadian drive function (phase delay) may be different in infancy and early childhood (hence why we set *b* = 1.0 in this work, based on previous findings (Webb et al 2024)), which would have an impact on sleep timing. This may explain why sleep bouts produced by our model had shifted start times compared to the real-life population of Mindell et al (2016) at the 1-year-old mark, and why the top insets in Figure 4 with a single bout of sleep per 24 hours have the sleep starting relatively late at night (after midnight in the instance of the top left insert). The results of the older age groups (1-, 3-, and 5-year-olds) suggest that *b* may have to shift closer to the adult value of *b* = 0.4 by the stage of more consolidated night sleep. In fact, changing *b* for the 5-year-olds had negligible impact on the *χ* and *μ*_ℎ*m*_ distribution, but did increase the area in parameter space yielding 24-h-entrained circadian rhythms. This parameter could also be explored in this framework using a probability model that includes a sleep timing outcome, such as sleep midpoint time as in Skeldon et al (2016), potentially revealing a joint, three-parameter shifting distribution with maturation. That said, the distribution of sleep midpoint time would be complicated for the infancy ages with several bouts given that sleep is distributed throughout the 24 h day, and we do not currently have access to sufficient data to test this.

Another limitation of this work is that the model only describes the sleep regulation system. Particularly in infancy, there are other influences that may affect sleep-wake behavior such as feeding drive, reaction to external noise, and inability to self soothe. The current model does not account for this, though we note that external stimuli could theoretically be incorporated within this model (Fulcher et al 2008). An additional shortcoming of the model fits is the excess density of 1-bout sleeps produced by the posterior compared to the empirical distribution, particularly for 1 y and 3 y of age (Figure 3e). This appears to be due to oversampling of the upper tail of the *log χ* distribution, where dynamics become insensitive to *χ* for large *χ*. Re-parameterising to sample in *log χ* is not sufficient to prevent the sampling dwelling excessively once it enters the one-bout regime of the *log χ* parameter space (Figure 3e, *log χ* ≳ 3.3). In future work, other parameterisations of *log χ* could be considered, or a different balance of chain and iterations in the sampling approach, or the need for additional sleep characteristics to constrain the model when it produces 1-bout consolidated sleep.

The data we fitted to also have some caveats. It is known that there are differences between sleep measures recorded through approaches such as actigraphy or polysomnography and parent-reported measures such as sleep diaries (Hall et al 2015, Mazza et al 2020). Parent-reported measures may inaccurately report sleep onset and offset, and hence misreport total sleep. Likewise, parents may not be aware of night wakings if the infant is able to soothe themselves back to sleep. Hence identifying bouts of sleep is inherently imprecise. We used empirical distributions based on several cohort studies and meta-analyses, including some that contain parent-reported measures. Potential biases between parent reported and automatic measures may shift the sleep characteristic distributions from the true population distributions, though we expect that the overall trends with age in both sleep characteristics and parameters will be robust.

## Conclusion

We estimated normative joint distributions for sleep model parameters for the heterogeneous sleep patterns observed across the early lifespan. This work sheds light on sleep homeostatic mechanisms underlying changing sleep patterns throughout infancy and childhood, in particular capturing both changes across time and the breadth of dynamics across the population. This work also establishes a proof of principle that population-level parameter estimation could be performed across the whole lifespan.

## Methods

### Sleep-Wake Switch Model

The Phillips and Robinson (2007) model is a neural mass model, where the sleep wake cycling behavior is produced by the mutual inhibition between the MA neuron group and the VLPO. Activity in the MA (*j* = *m*) and VLPO (*j* = *v*) neuron populations obeys

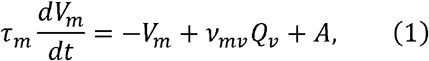

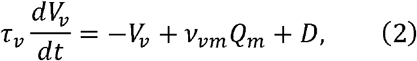

where *V_j_* is the mean cell body potential of neuronal population *j* = *m*, *v*, with membrane time constant *τ_j_* , and *Q_j_* is its mean firing rate, and *A* is a constant excitatory input from the orexin and Ach group (Figure 1). Firing rates are a sigmoidal function of cell body potentials, given by

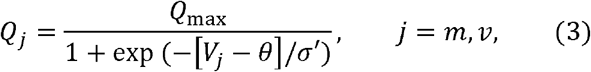

where *θ* is the firing threshold, *σ*′ can be interpreted as a spread in the threshold or a distribution in activity levels around the mean, and *Q_max_* is the maximum firing rate.

The term D in Equation 2 is the sleep-promoting drive to the VLPO, and is the sum of the homeostatic drive (H), circadian process (C), and a constant background input to the VLPO (D_0_):

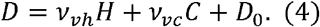

The *ν_ab_* terms represent the strength of the input from the neuron population/drive *b* into neuron population *a*. For example, *ν_vc_* is the strength of the circadian input into the VLPO (part of the sleep drive *D*).

The homeostatic process *H*, the process by which sleep inducing chemicals such as adenosine build up in the brain (particularly the basal forebrain) during wakefulness and clear during sleep, obeys

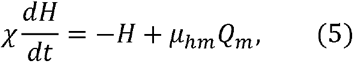

where *χ* is the characteristic clearance time of the somnogens, and *μ*_ℎ*m*_ is the rate of increase in the level of somnogens from the wake promoting MA group.

The circadian process *C* is defined by

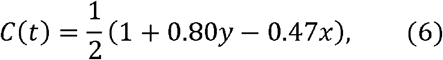

modelled on a forced van der Pol oscillator (light entrainment of the circadian process) defined by

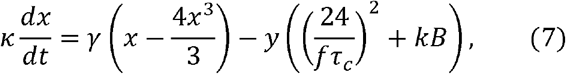

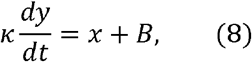

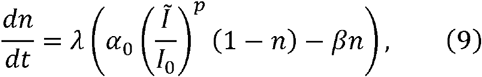

where

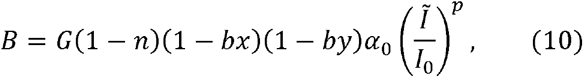

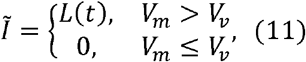

and where *L*(*t*) defines the solar curve, obeying

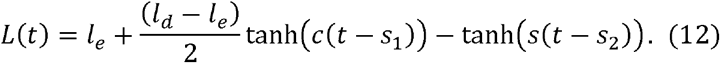

The van der Pol oscillator variables are *x* and *y* (Equation 7 and 8), with *n* being the fraction of activated photoreceptors (Equation 9).

In Equation 9, the photoreceptor rate of saturation is governed by 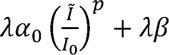 and rate of decay in the absence of light input by *λb*. The stiffness of the oscillator is *γ* (Equation 7), with *τ_c_* being the intrinsic period, *f* a small correction to the period, and *k* the dependence of the intrinsic period on the light intensity. The circadian oscillator has varying sensitivity to light in different phases (Skeldon et al 2017). The term *B* describes the forcing resulting from the photoreceptors being acted upon by the light intensity *I*, with the sensitivity to phase being determined in Equation 9 by (1 − *bx*)(1 − *by*). In the solar curve in Equation 12, *l_e_* is the minimum light level in the evening and *l_d_* is the maximum light level during the day. The term *c* sets the steepness of the solar curve, with the change from *l_e_* to *l_d_* occurring around the time *s*_1_, and the change from *l_d_* to *l_e_* occurring around the time *s*_2_. The value for *l_e_* is set to 0 for this infant/young child case. As identified in previous work (Webb et al 2024), we set *b* = 1.0 as this value better aligns with infant and young child sleep. This parameter changes the response to the solar curve, and its effects on sleep timing can be seen in Appendix S4. Wake is defined as when *V_m_* > *V_v_* . Parameter values were taken from Skeldon et al (2017) unless otherwise noted, and are given in Table 1.

### Bayesian model

Due to the large variability in sleep characteristics observed in infancy, it is unrealistic to expect a single set of parameter values (*φ_age_* ) to generate the diversity of sleep characteristics at a single age. Instead, estimating parameters with some estimate of spread is more useful to describe the distribution of parameter values able to generate the observed population-level diversity of sleep characteristics. We used Bayesian analysis to estimate the joint distribution *P*(*ψ_i_*|*Π_i_*), where *ψ_i_* is the set of parameters in the Phillips and Robinson sleep model at a particular age *i*, and *Π_i_* is a collection of probability density functions describing the distribution of sleep characteristics at a given age *i*. We used flat priors on the parameters, taking into account range constraints based on known physiological bounds (Phillips & Robinson 2008).

We did not have access to a dataset of sleep behavior representing a whole population of a single age of infancy, so instead we used reported summary statistics from meta-analyses and cohort studies (Dias et al 2018, Iglowstein et al 2003, Jenni & Carskadon 2012, Sadeh et al 2009, Staton et al 2020) (Appendix S1) to construct probability distributions (Figure 2, Table 2). Specifically, we constructed sleep behavior characteristic probability distribution (*Π_i_*) models at seven different ages: 1, 3, 6, and 9 months old, and 1, 3, and 5 years old. We focused on explaining two key features of infant and young child sleep: the increased amount of sleep, and its polyphasic nature (day time naps and night wakings). Additionally, our measures had to be objectively calculable from the sleep-wake model time series output. Hence, we aimed to fit two key sleep behavior characteristics: the average amount of sleep per day and the average number of sleep bouts per day (an objective summary of both naps and night-waking). We assumed that the distribution of average amount of sleep per 24 hours at any given age was normally distributed:

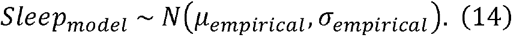

The number of bouts of sleep is an integer within a day but not necessarily when averaged across many days; for example, an infant in a behavioral regime of 5 bouts of sleep per 24 hours might have 5 bouts of sleep most days but on some days have 4 or 6 bouts of sleep. We assumed a mixed Gaussian distribution of number of sleep bouts per 24 hours for each age,

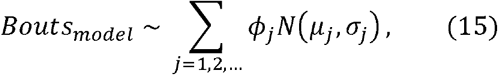

with the individual Gaussians (weighted by *φ_j_*) centered on integers (*μ_j_* ∈ 1,2, …) and the standard deviation (SD) of each Gaussian proportional to the mean (*σ_i_* = *μ_i_*/10). Proportionality of mean to SD was chosen as it was assumed that the variation in number of bouts would be larger when bouts per day is higher, and sensitivity analysis for the magnitude of this proportional relationship is given in Appendix S5. The mixture weightings of the Gaussians (*φ_j_*) were chosen to produce summary statistics found in the literature (Table 2, Supplementary Table S1). For example, the *φ_j_* used for 5 years of age were selected to have P(X≥1)≈7%, to match the reported statistic in the literature of only 7% of 5-year-olds still having a daily nap (Staton et al 2020).

A large range of *χ* can produce monophasic sleep, and as *χ* decreases the sleep breaks into increasing numbers of bouts. As small changes in *χ* when *χ* is small can have large changes in the number of bouts of sleep, we reparametrized the model to use *χ* on a natural log scale, so as to have a more balanced parameter space for sampling across numbers of bouts.

### Implementation

To perform the Bayesian analysis, we implemented the Phillips and Robinson model in the probabilistic programming language Stan (using cmdstan via cmdstanr (Stan Development Team 2020); code available: https://github.com/LockyWebb/SleepStanModels). The solver function ode_bdf (Stan Development Team 2020), a backwards differentiation formula (BDF) method solver, was used to solve the model at fixed time steps of 18 seconds. Stan uses an adaptive Hamiltonian Monte-Carlo (HMC) sampling with a No-U-Turn Sampler (NUTS) (Stan Development Team 2019).

### Bayesian Model

Our modeling approach used *μ*_ℎ*m*_ to change the amount of sleep per day, as per Skeldon et al (2016) and other posited theories in the literature (Gibson et al 2012, Jenni & LeBourgeois 2006, Spencer & Riggins 2022). Previous work constrained the range of *μ*_ℎ*m*_ to 2.1 nM s to 7.2 nM s, based on the healthy sleep amount for adults of 6 to 10 hours (Phillips & Robinson 2008). Extending that to a possible 18 hours of sleep per day for infants gives an upper bound of ∼16 nM s. Hence, we used a prior of *μ*_ℎ*m*_∼*U*(2.1,16.0), where *U*(*a*, *b*) denotes the uniform distribution on the interval [*a*, *b*]. We used a uniform prior for *χ* on the natural logarithmic scale of *log χ* ∼*U*(1.5,4.0) (range of *χ* 4.5 to 54.6 h). This range of *χ* includes the previously used adult value of 45 h. The use of *μ*_ℎ*m*_ aligns with modeling done by Skeldon et al (2016).

### Model execution and sleep summary statistics

We ran the Bayesian model for each age using cmdtsan (Stan Development Team 2020). Each instance of the model was run sampled on 20 parallel chains with 500 warm-up iterations and 500 sampling iterations, using a max tree depth of 7 and adaption delta of 0.5 (Stan Development Team 2019). Choices for chain and iteration numbers were made to balance the desire for a large number samples (20 × 500 = 10000) and computational time and resources. HMC parameters (tree depth and adaption delta) were chosen over default values to improve computational times, after confirming in small test runs that it didn’t impact the resulting posteriors.

The sleep model was run for 35 days in each iteration. The first 21 days of this simulation was considered a warm up for the sleep model to remove any transient behavior, with the last 14 days summarized by the sleep behavior characteristics of average amount of sleep per 24 hours and average number of bouts per 24 hours.

### Assessing observed circadian pacemaker period

To determine whether the model was entrained to the 24-hour day, the observed period (*τ_obs_*) of the circadian pacemaker (the modified forced van der Pol oscillator, Equations 7 and 8) was calculated by,

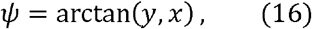

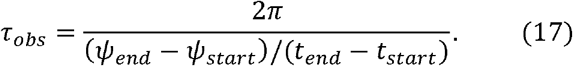

## Code availability

Code for the STAN models used is available at https://github.com/LockyWebb/PR_Sleep_Stan_Models.

## Data availability

All data used are based on summary statistics from cited research and summarized in supplementary material (Supplementary Table S1).

## Supplementary Material

**Appendix S1 – Data sources**

**Supplementary Table S1:** Published summary statistics of sleep characteristics used to construct the empirical distributions of sleep amount and bout number at each age modelled.

**Appendix S2 – Bayesian Model: *D*_0_ − *χ***

**Supplementary Figure S1:** Age-specific parameter estimation in the *D*_0_ − *χ* model.

**Supplementary Figure S2:** Contours from a kernel density estimate containing 95% of the joint posterior sampled points of *D*_0_ and log *χ* posteriors at each age.

**Appendix S3 – Observed circadian period in sleep patterns**

**Supplementary Figure S3**: The observed circadian period for different sleep patterns.

**Supplementary Figure S4**: The observed circadian period for in the *μ*_ℎ*m*_ − log *χ* space, combining the parameter distributions for each age.

**Supplementary Figure S5**: The observed circadian period for in the *D*_0_ − log *χ* space, combining the parameter distributions for each age.

**Appendix S4 – Effect of parameter *b***

**Supplementary Figure S6**: Effect on sleep patterns of parameter b being equal to either 1.0 or the nominal adult value 0.4.

**Supplementary Figure S7**: Effect on 5-year-old *μ*_ℎ*m*_− *χ* distribution when *b* is changed.

**Supplementary Figure S8**: The observed circadian period for in the *μ*_ℎ*m*_ − log *χ* space in the 5-year-old case.

**Appendix S5 – Sensitivity analysis of BPD to the Mixed Gaussian**

**Supplementary Figure S9**: Results from the *D*_0_ − *χ* model with different empirical distribution (Gaussians with smaller variances) for average number of bouts.

