## Supplementary Material for "Inferring population-level physiologically based model parameters for sleep in infancy and young childhood"

1) Brain Modelling Group, QIMR Berghofer Medical Research Institute, Herston, Brisbane, QLD 4006, Australia
2) Faculty of Medicine, University of Queensland, QLD, Australia
3) Turner Institute for Brain and Mental Health, School of Psychological Sciences, Monash University, Clayton, VIC, Australia

**Appendix S1 – Data sources**

**Appendix S2 – Bayesian Model:** $\boldsymbol{D}_{\boldsymbol{0}}\boldsymbol{-\chi}$

**Appendix S3 – Observed circadian period in sleep patterns**

**Appendix S4 – Effect of parameter** $\boldsymbol{b}$

**Appendix S5 – Sensitivity analysis of BPD to the Mixed Gaussian**

**Appendix S1**

**Data sources**

Supplementary Table S1 describes the literary sources behind the population distribution values used.

**Supplementary Table S1:** Published summary statistics of sleep characteristics used to construct the empirical distributions of sleep amount and bout number at each age modelled.

|  | **Sleep (h)** | | **Source** | **Bout** | | **Source**  Using justification of:  $bouts\approx naps+night wakings-1$ |
| --- | --- | --- | --- | --- | --- | --- |
| **Age** | **Average (**$\boldsymbol{\mu}$**)** | **SD (**$\boldsymbol{\sigma}$**)** |  | **Average (**$\boldsymbol{\mu}$**)** | **SD (**$\boldsymbol{\sigma}$**)** |  |
| 1m | 14.4 | 3.00 | Iglowstein et al (2003) average $\approx14.3$ h, Galland et al (2012) average $=14.6$ h, Sadeh et al (2009) average $=14.3$ h. max SD from Sadeh et al (2009) | 6.005 | 1.4542 | Galland et al (2012): mean 3.1 naps, with minimum of 1.2 and maximum of 5.0, and mean night wakings of 1.7, minimum of 0 to maximum of 3.4. Sadeh et al (2009) mean (SD) of 3.59 (1.18) naps and 1.89 (1.09) night wakings. |
| 3m | 13.8 | 2.14 | Iglowstein et al (2003) average $\approx14.5$ h, Galland et al (2012) average $= 13.6$ h, Sadeh et al (2009)average $=13.3$ h. Max SD from Iglowstein et al (2003). | 5.207 | 1.2154 | Galland et al (2012): mean 3.1 naps, with minimum of 1.2 and maximum of 5.0, and mean night wakings of 0.8, minimum of 0 to maximum of 3.0. Sadeh et al (2009) mean (SD) of 2.93 (0.83) naps and 1.24 (1.19) night wakings. |
| 6m | 13.0 | 2.10 | Sadeh et al (2009) average =12.9 h, Galland et al (2012) average = 12.9 h, Iglowstein et al (2003) average = 14 h. from Dias et al (2018), Konrad et al (2016) average = 12.6 h, Galland et al (2016) average = 10.4 from actigraphy and average = 12.8 from sleep diary. Blair et al (2012) average = 13.2 h with wide age bin. Average from all sources is 13.0 h. Max SD is from Galland et al (2012). | 4.675 | 1.324 | Galland et al (2012): mean 2.2 naps, with minimum of 0.9 and maximum of 3.5, and mean night wakings of 0.8, minimum of 0 to maximum of 3.0. Sadeh et al (2009) mean (SD) of 2.42 (0.75) naps and 1.25 (1.20) night wakings. |
| 9m | 12.8 | 1.74 | Iglowstein et al (2003) average $\approx$ 13.9 h, Galland et al (2012) average = 12.6 h, Sadeh et al (2009) average = 12.8 h. Using Sadeh et al (2009) as middle of average values and well reported trend between 6 months and 12 months.  Max SD from Sadeh et al (2009). | 4.135 | 1.166 | Galland et al (2012): mean 2.2 naps, with minimum of 0.9 and maximum of 3.5, and mean night wakings of 1.1, minimum of 0 to maximum of 3.1. Sadeh et al (2009) mean (SD) of 2.02 (0.52) naps and 1.16 (1.17) night wakings. |
| 12m | 12.8 | 1.59 | Iglowstein et al (2003) average =13.9 h, Galland et al (2012) average = 12.9 h, Sadeh et al (2009) average = 12.8 h. Dias (2018) reports values form three studies, 11 h (diary, 12.25 h with actigraphy), 13 h, and 12.2 h.  Max SD from Sadeh et al (2009). | 3.225 | 0.968 | Galland et al (2012): mean 1.2 naps, with minimum of 0.4 and maximum of 2.1, and mean night wakings of 0.7, minimum of 0 to maximum of 2.5. Sadeh et al (2009) mean (SD) of 1.53 (0.55) naps and 0.93 (1.18) night wakings. Gibson et al (2012) states 4 sleep periods. |
| 36m | 12.0 | 1.54 | Iglowstein et al (2003) average $\approx12.5$ h, Galland et al (2012) average = 12.0 h, Sadeh et al (2009) average = 11.9 h. Max SD from Sadeh et al (2009). | 2.73 | 0.959 | Sadeh et al (2009) has 24 to 36-month-olds with mean (SD) naps of 0.92 (0.37) and 0.82 (0.92) night wakings. |
| 60m | 11.4 | 1.22 | Iglowstein et al (2003) average $\approx$ 11.4 h, Galland et al (2012) average = 11.5 h. Max SD from Galland et al (2012). | 1.07 | 0.2778 | Staton et al (2020) estimated 93% of 5-year-olds had ceased napping using low risk studies. Assuming no night wakings by that age. |

**Appendix S2**

Bayesian model: $D_{0}-\chi$ results

We additionally explored the results of the model where $\boldsymbol{\psi}_{\boldsymbol{i}}$ for each age $i$ contains $D_{0}$, the constant inhibition to the VLPO (directly influencing the amount of sleep), and $\chi$ (the homeostatic timescale). This combination of parameters was previously used to explain the diversity in sleep behavior across different mammalian species (Phillips et al 2010).

The $D_{0}-\chi$ model had a fixed $\mu_{hm}=4.5$ nM s as per previous modeling work in adults, with priors $D_{0}\sim U\left( -11,-1.0 \right)$ and $\log\chi\sim U\left( 1.5,4.0 \right)$ ($\chi$ ranging from 4.5 to 54.6 h, same prior as the main model that used $\mu_{hm}$and $\chi$). These ranges of $D_{0}$ and $\chi$were chosen as they covered the full range of sleep behavior from adulthood down to infancy.

This model yielded similar trends across ages to the first model (which had $\mu_{hm}$ in place of $D_{0}$), with the distributions for $D_{0}$ at younger ages being wide and centered at higher values, and lower-centered distributions for the older ages (Supplementary Figure S1a). The distributions for $\log\chi$ shifted higher with age (Supplementary Figure S1b), and formed clusters (bumps in the violin plots) that align with different integer numbers of bouts, reflecting a step-like relationship between number of bouts and $\chi$ for a fixed $\mu_{hm}$. The sharp change at $\log\chi\sim$3.2 for age 1 y and up reflects the change from multiple bouts of sleep per day to a single consolidated bout of sleep per day. The trends in $D_{0}$ (higher values yield more sleep) and $\log\chi$ (higher values yield more fragmented sleep) are consistent with prior work examining sleep patterns across species (Phillips et al 2010).

The distribution of $D_{0}$ for early ages hits the assumed upper boundary of $D_{0}=-1.0$ (Supplementary Figure S1a), beyond which total sleep per day approaches 24 h. This is likely due to the upper tail of the empirical density function ($\Pi_{i}$) of sleep per 24 hours having non-zero weight for over 20 hours of sleep per 24 hours for the younger ages $(i=1 m,3 m)$. At this $D_{0}$ upper bound the maximum sleep per 24 hours is over 20.3 hours, which could reasonably be considered the limit of not-severely-pathological sleep at age 1 month. Similar to the results in the first model, the posterior distributions of $\log\chi$ also hit the upper $\log\chi$ boundary in the older ages where sleep is monophasic (Supplementary Figure S1b).

Visual comparison of the distributions of sleep characteristics generated by the model posteriors shows excellent agreement with the empirical estimates (Supplementary Figure S1d). While the model could produce non-integer average numbers of bouts per day (33% were non-integer), the level of discreteness of the average number of bouts per day exceeded that of the assumed population distribution (Supplementary Figure S1e).

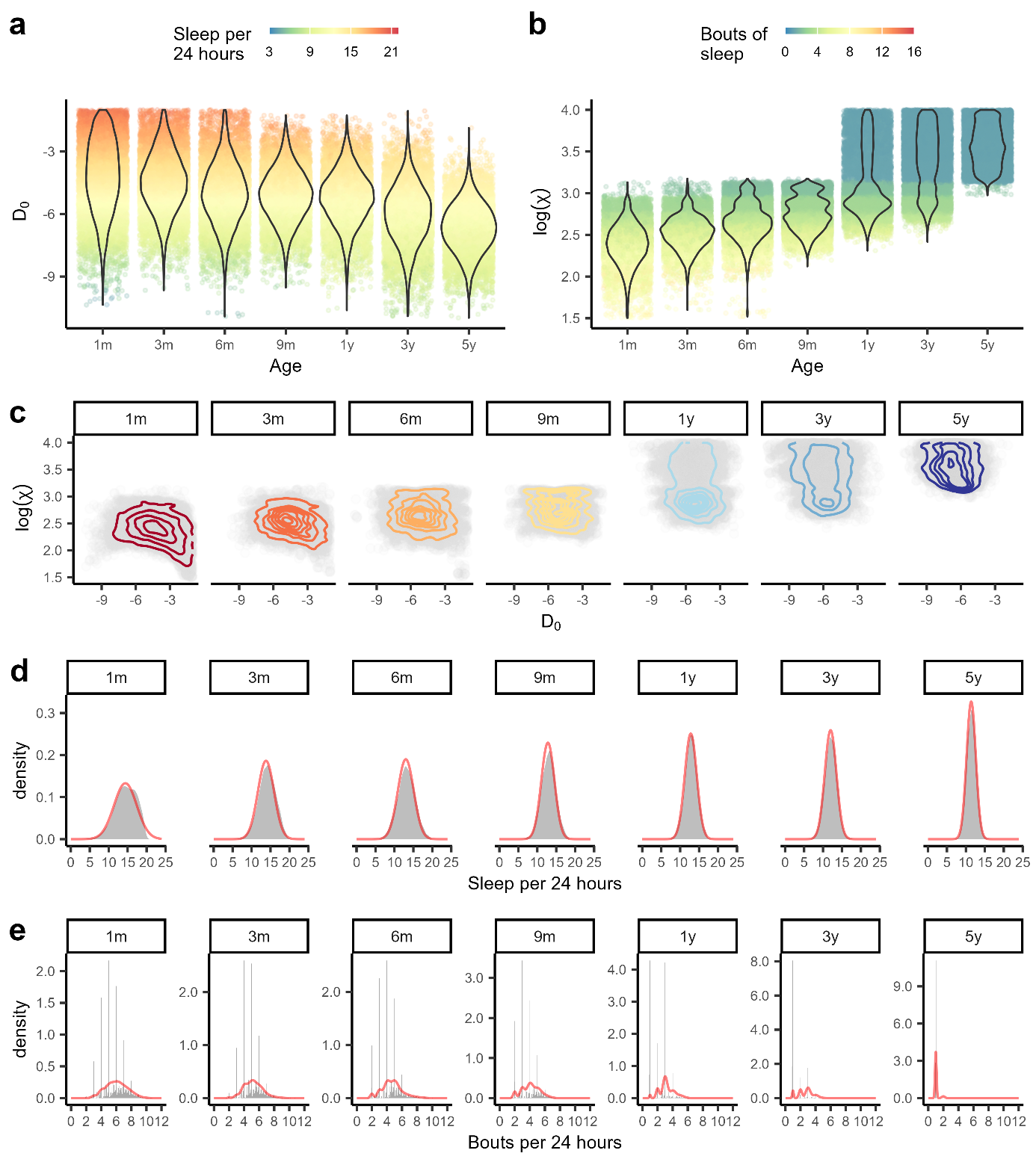

**Supplementary Figure S1:** Age-specific parameter estimation in the $D_{0}-\chi$ model. **a)** Posterior distributions (violins) for each age for $D_{0}$, with points colored by the corresponding average sleep per 24 hours. **b)** Posterior distributions (violins) for each age for $\log\chi$, with points colored by the corresponding number of bouts of sleep. **c)** The joint posterior distribution (gray) for each age with density contours shown at density levels 0.05,0.01,0.15,.... **d)** Distributions by age for the amount of sleep per 24 hours arising from the parameter posteriors (gray) superimposed with the empirical density curves (red). **e)** Distributions for the average number of bouts per 24 hours.

The centers of the density contours decrease in $D_{0}$ and increase in $\chi$ with age (Supplementary Figure S2). As in Figure 4, the outer contours in Supplementary Figure S2 show that there is wide population variation, with the distributions contracting with increasing age. There is again a small overlap between 5-year-olds and 9-month-olds, indicating the model parameters of the most consolidated 9-month-olds are similar to those of the small amount of 5-year-olds exhibiting polyphasic sleep.

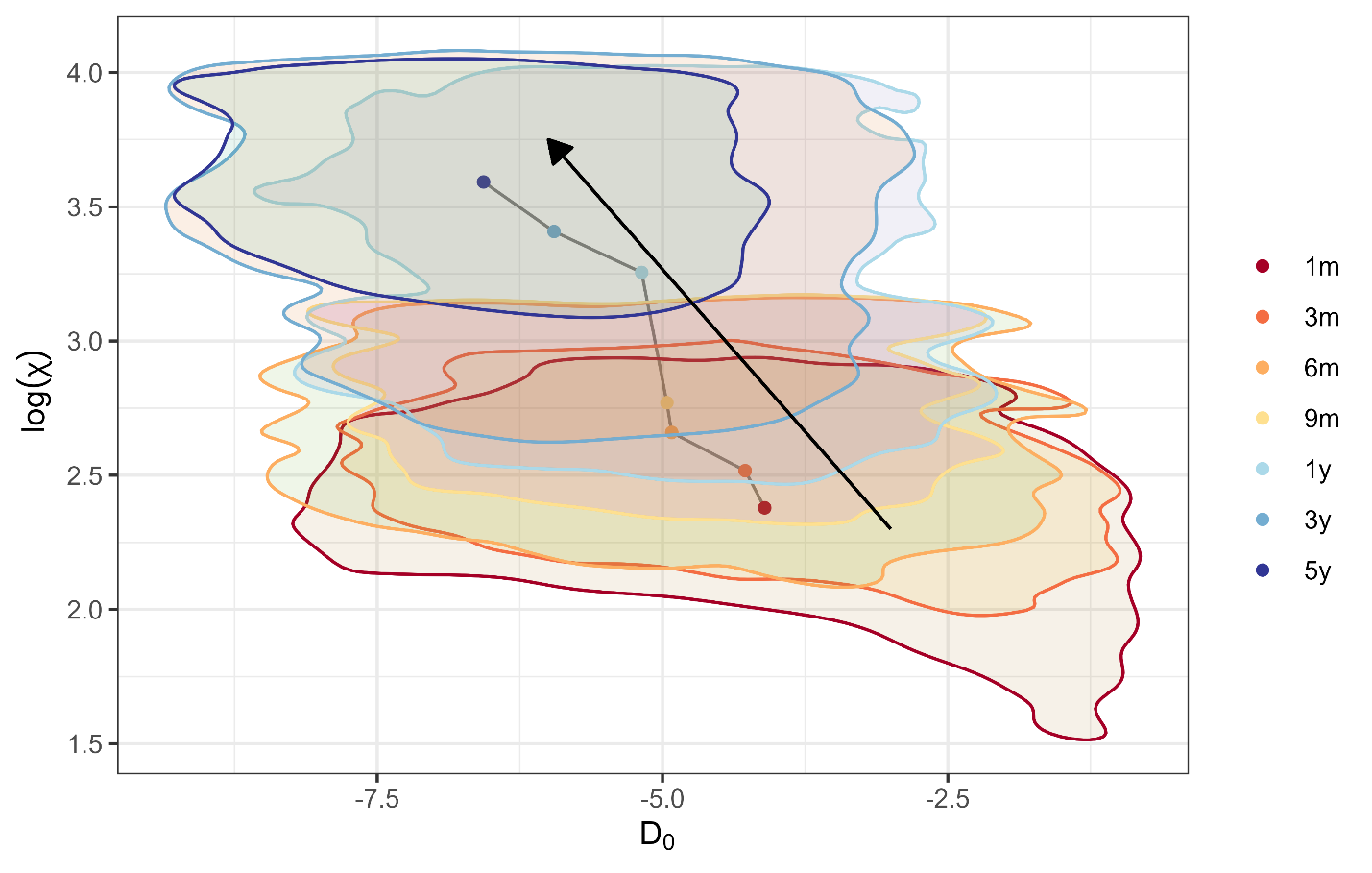

**Supplementary Figure S2:** Contours from a kernel density estimate containing 95% of the joint posterior sampled points of $D_{0}$ and $\log\chi$ posteriors at each age. Centrers of distributions are shown with dots, colored by age; arrow denotes the direction of increasing age. Insets show examples of modelled sleep behavior at the parameters indicated by black dots, with light blue bars signifying bouts of sleep.

Overall, the $D_{0}-\chi$ model appears to yield a better fit to the empirical population sleep characteristic probabilities ($\Pi$) than the $\mu_{hm}-\chi$ model (Figure 3), mainly evident by the discrepancy between $\Pi_{i}$ and $\psi_{i}$ for $i=$ 1 month, 3 month, and 6 month in the $\mu_{hm}-\chi$ model for the average amount of sleep per 24 hours (Figure 3d). This may be due to the greater independence of $D_{0}$ and $\chi$ than $\mu_{hm}$ and $\chi$.

**Appendix S3**

**Observed circadian period in sleep patterns**

The distribution of sleep patterns for the 1-year-old results in Figure 7 included a small proportion of sleep episodes that lasted 10-12 hours and began at any point throughout the 24 hours of the day. Supplementary Figures S3a and S3b show, for the $\mu_{hm}-\chi$ model of the main paper and the $D_{0}-\chi$ model of Appendix S2, respectively, that the observed circadian period is much higher than 24 h (or even the natural period of approximately 24.2 h). The sleep episodes that last 12 hours and begin between 6am and 12pm have the highest period. The sleep bouts from patterns with observed circadian period less than 24.2 hours are shown in Supplementary Figures S3c and S3d, where the sleep start clock-time distribution of sleep episodes over 10 hours is much narrower, now confined to the afternoon and evening. Only a small proportion of model produced sleep patterns have a period close to 24 h (<24.1 h and >23.9 h), with the smallest proportion for the 1 month of age results (9% and 2% for the $\mu_{hm}-\chi$ and $D_{0}-\chi$ models, respectively) and the largest proportion for the 1-year-old results (22% and 16% for the $\mu_{hm}-\chi$ and $D_{0}-\chi$ models, respectively). It should be remembered that we are not modeling any external influences on the sleep-wake circuitry beyond light.

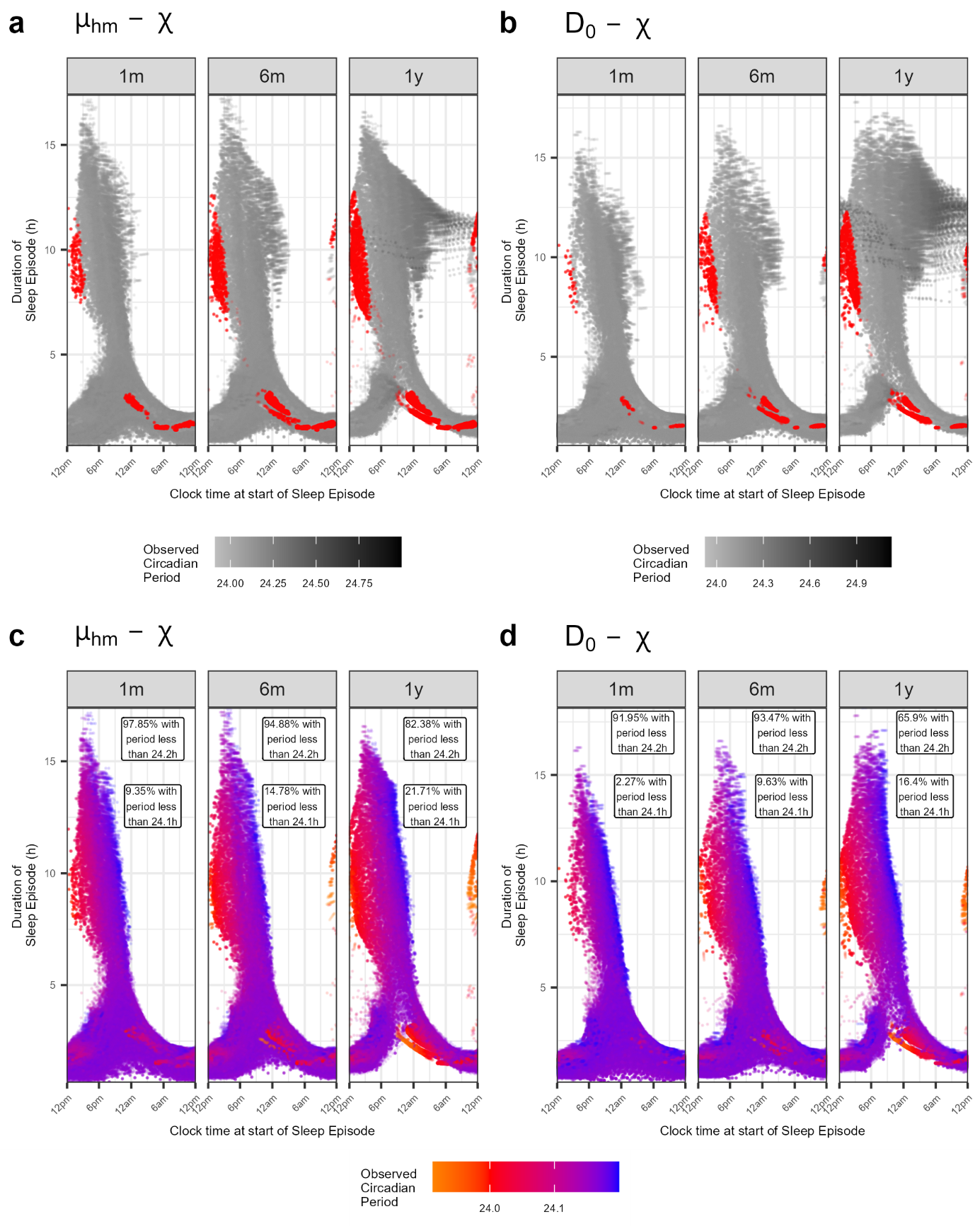

**Supplementary Figure S3**: The observed circadian period for different sleep patterns. The sleep episode length and clock start time for **a)** the $\mu_{hm}-\chi$ model and **b)** the $D_{0}-\chi$ model, as per Figure 7, with the points colored by the observed circadian period (observed periods between 23.95 and 24.05 h highlighted in red). The sleep patterns when the observed circadian period is less than the inherent 24.2 h natural rhythm are shown in **c)** for $\mu_{hm}-\chi$ and in **d)** for $D_{0}-\chi$. The proportion of points in **a)** and **b)** that have a period less than 24.2 h and 24.1 h are displayed in **c)** and **d)**.

When inspecting the parameter space of the two models using all points in all posteriors (all ages combined, Supplementary Figure S4 and Supplementary Figure S5 for $\mu_{hm}-\chi$ and $D_{0}-\chi$, respectively) it can be seen that the highest observed circadian period is produced by high $\log\chi$ and low $\mu_{hm}$ or $D_{0}$ (Supplementary Figure S4a and Supplementary Figure S5a). Those points are in the single bout of sleep per day parameter space (Supplementary Figure S4b and Supplementary Figure S5b), and would correspond with the points at the edge of the parameter posteriors in Figure 3a and b, and Supplementary Figure S1a and b. Focusing just on model-produced sleep that has a observed circadian period of close to 24 h (<24.1 h and >23.9 h) reveals small clusters of points, mostly still above 24 h (Supplementary Figure S4c and Supplementary Figure S5c), with each cluster generally a different whole number of bouts of sleep per day (Supplementary Figure S4d and Supplementary Figure S5d). These clusters approximately align with the trajectory of the centers of the contours in Figure 4 and Supplementary Figure S2.

It is possible that parameters producing un-entrained and unrealistic sleep could reflect the underlying mechanisms in some infants/children, and that external influences such as parents could entrain the sleep patterns more realistic behaviors. However, these results highlight that the unrealistic sleep behavior at the boundaries of the parameter ranges (in particular the upper boundary of the $\log\chi$ posterior distributions) are likely not reflective of true infant/child sleep, and that future work with more sleep measures (such as observed circadian period) in the probability model, and changes in other sleep model parameters (such as $b$, see Appendix S4), is needed to ‘tighten’ the posteriors.

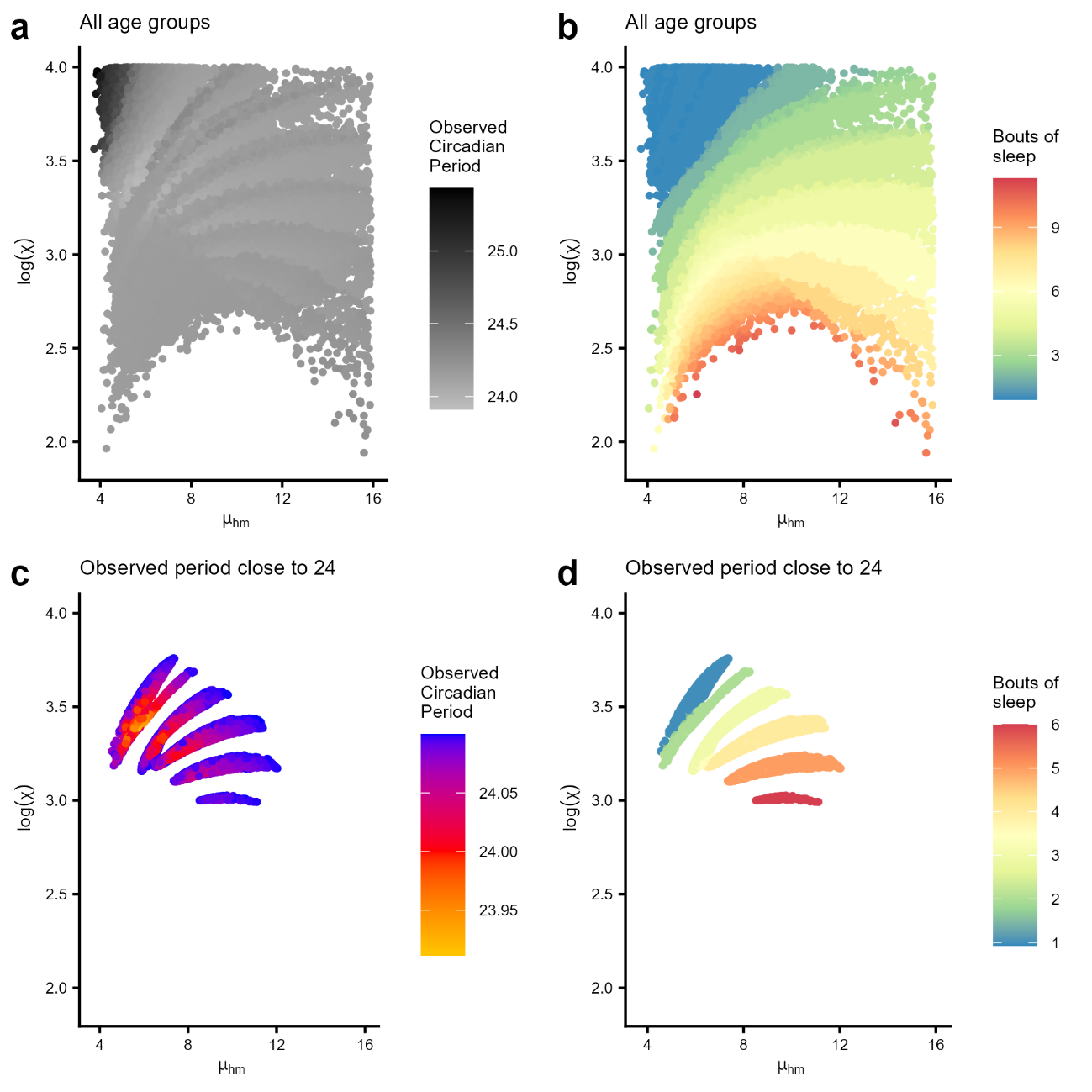

**Supplementary Figure S4**: The observed circadian period for in the $\mu_{hm}-\log\chi$ space, combining the parameter distributions for each age. **a)** The points from each posterior colored by the observed circadian period, showing the points with the highest period focused at low $\mu_{hm}$ and high $\log\chi$. The correspoding number of bouts per day in this parameter space is shown in **b)**, where divides between whole numbers of bouts (with a small amount of non-whole number between) is evident. Focusing on just the parameter combinations with period close to 24 h (<24.1 h and >23.9 h) gives small clusters of points in the space, colored by **c)** observed circadian period, and **d)** number of bouts of sleep per day.

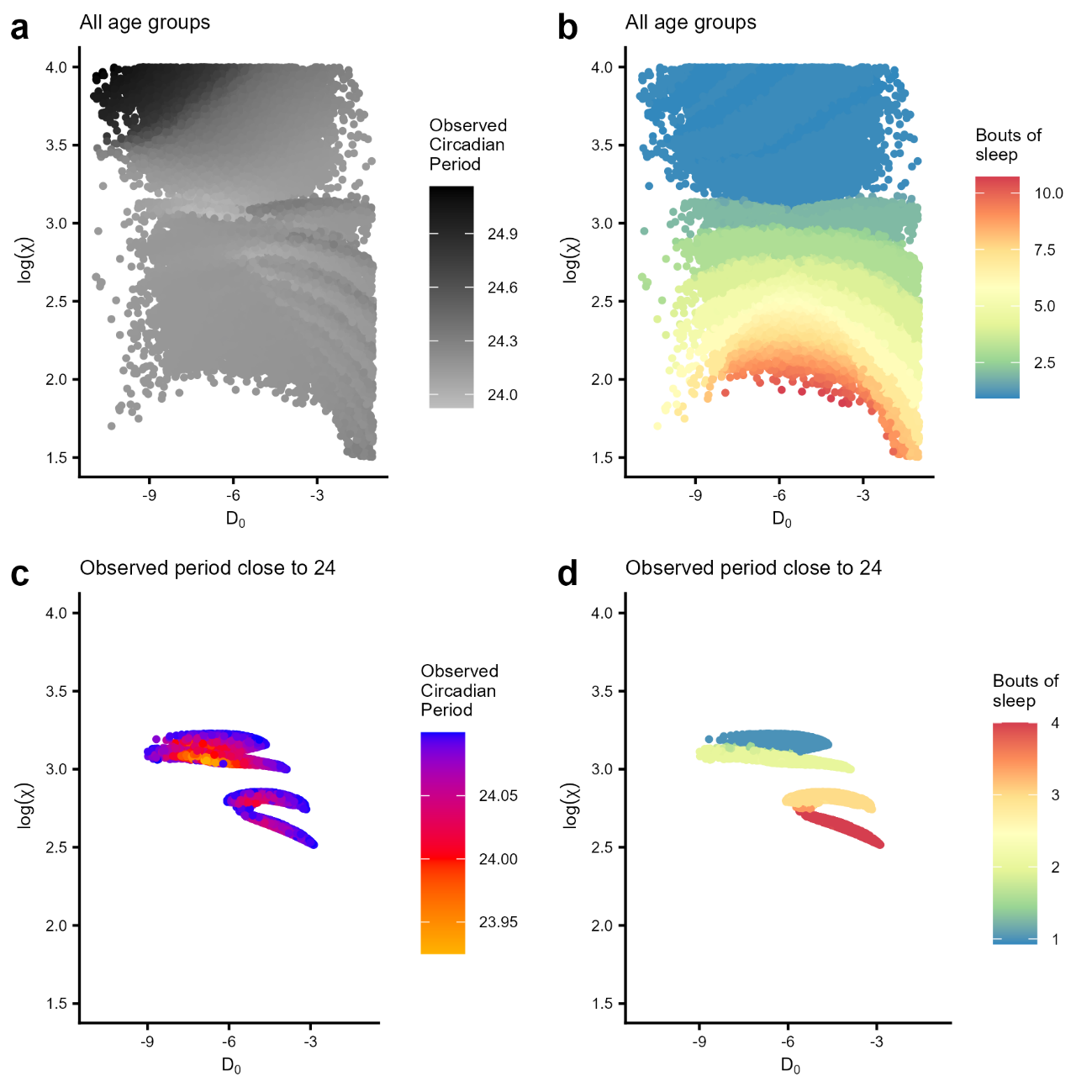

**Supplementary Figure S5**: The observed circadian period for in the $D_{0}-\log\chi$ space, combining the parameter distributions for each age. **a)** The points from each posterior colored by the observed circadian period, showing the points with the highest period focused at low $D_{0}$ and high $\log\chi$. The correspoding number of bouts per day in this parameter space is shown in **b)**, where divides between whole numbers of bouts (with a small amount of non-whole number between) is evident. Focusing on just the parameter combinations with period close to 24 h (<24.1 h and >23.9 h) gives small clusters of points in the space, colored by **c)** observed circadian period, and **d)** number of bouts of sleep per day.

**Appendix S4**

**Effect of parameter** $\boldsymbol{b}$

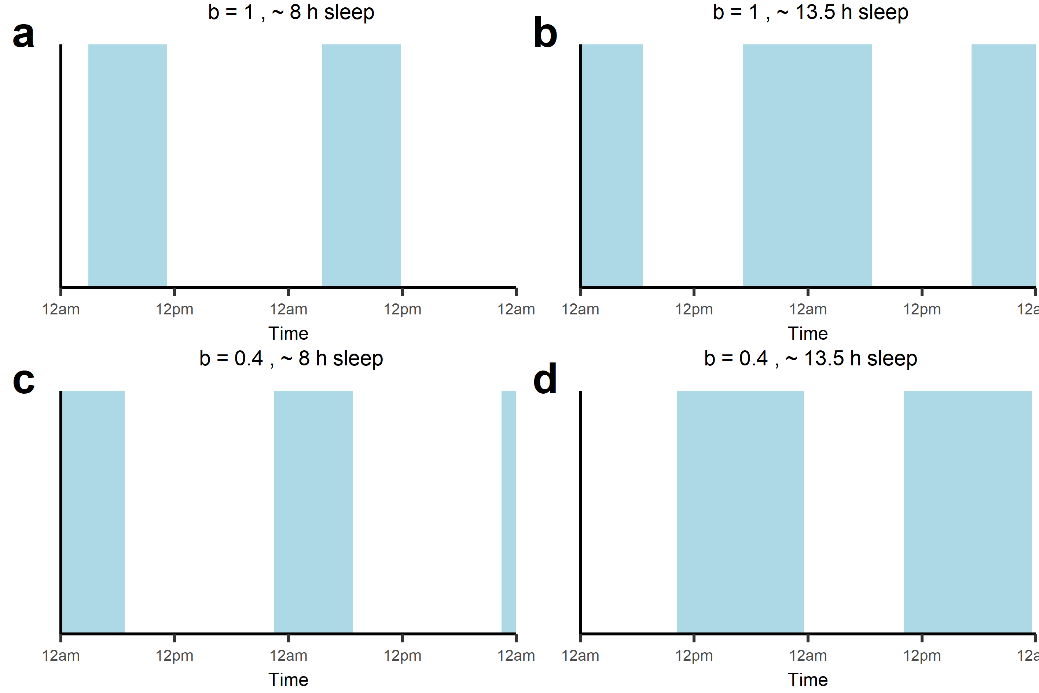

**Supplementary Figure S6**: Effect on sleep patterns of parameter b being equal to either 1.0 or the nominal adult value 0.4. **(a)** Sleep patterns with $b=1.0$ for ~8 hours sleep. **(b)** $b=1.0$ for ~13.5 hours of sleep). **(c, d)** Sleep patterns for with $b=0.4$ for ~8 hours sleep and ~13.5 hours of sleep.

The parameter $b$, the phase response of the circadian pacemaker to the solar curve, was set to $b=1.0$ for all results in this paper. In the parameter space with a single sleep bout per day, a $b$ value of 1.0 (Supplementary Figure S6a and b) results in sleep with unrealistic timing for the shorter sleep bout (sleeping until midday, falling asleep after midnight in the 8 hour of sleep example etc.), but realistic for the longer sleep bout. While when $b=0.4$ (Supplementary Figure S6c and d), the timing of the shorter sleep bout is much more realistic but the longer sleep bout has unrealistic timing, sleeping from just before midday to almost midnight. We set $b$ to 1.0 as it is needed for the infant case (Webb et al 2024) and didn’t allow it to vary as part of $\psi_{i}$ as we were not fitting to any sleep timing outcomes.

While not fit to sleep timing outcomes, when the parameter $b$ was changed to different fixed values for the 5-year-old case (the most likely of the ages considered to need a different value for $b$), the distribution of $\mu_{hm}$ and $\chi$ has negligible change (Supplementary Figure S7).

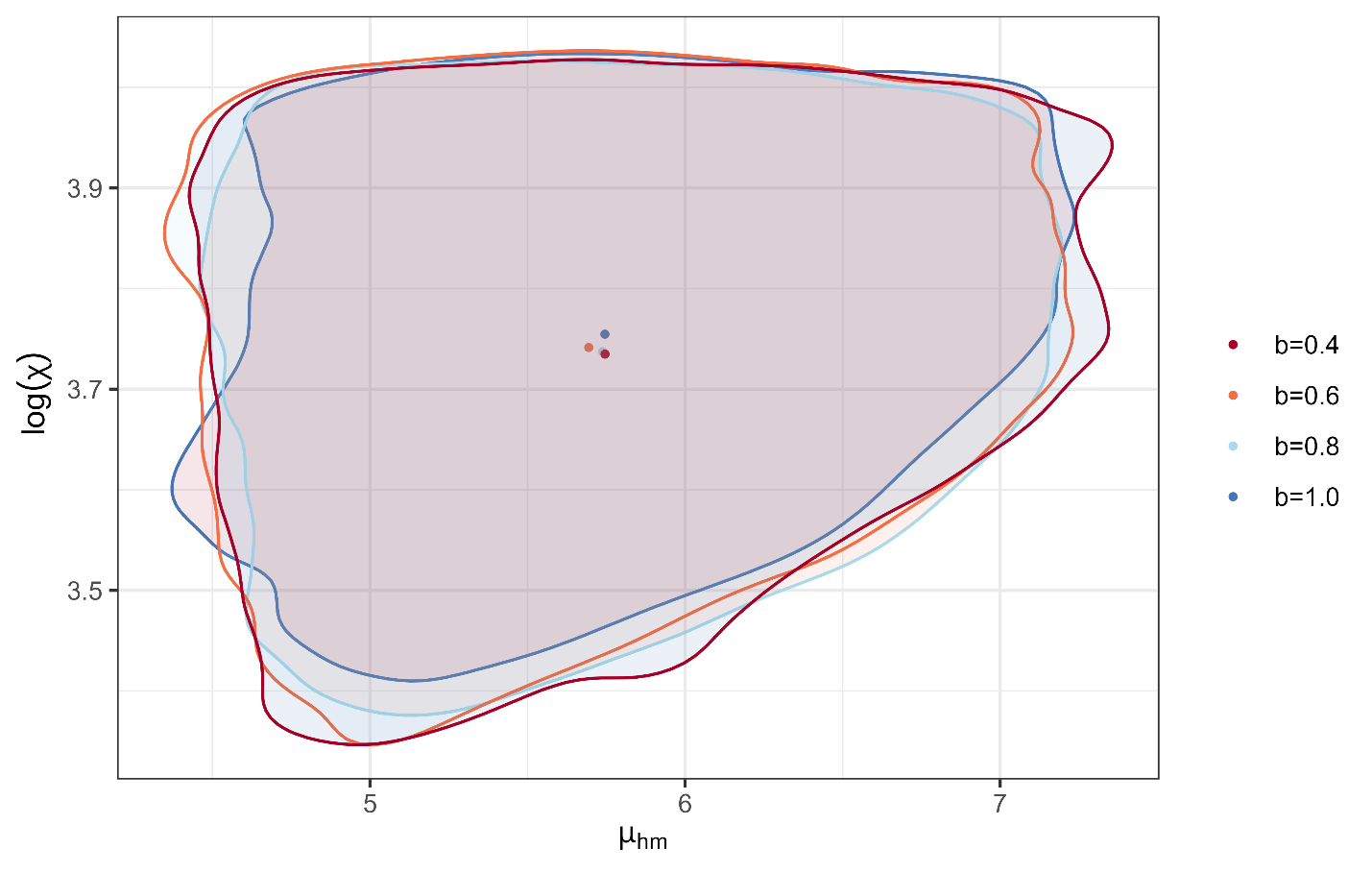

**Supplementary Figure S7**: Effect on 5-year-old $\mu_{hm}-\chi$ distribution when $b$ is changed. Contours from a kernel density estimate containing 95% of the joint posterior sampled points of $\mu_{hm}$ and $\log\chi$ posteriors for each value of $b$. Centers of distributions are shown with dots, colored by value of $b$.

When exploring the observed circadian period in these $\mu_{hm}-\chi$ distributions, it was revealed that the observed circadian period was generally closer to 24 hours in the cases of smaller $b$ (Supplementary Figure S8). Additionally, the $\mu_{hm}-\chi$parameter space that allowed observed circadian periods close to 24 hours was larger for $b=0.4$, mostly made up of parameter combinations that produce 1 bour of sleep per day.

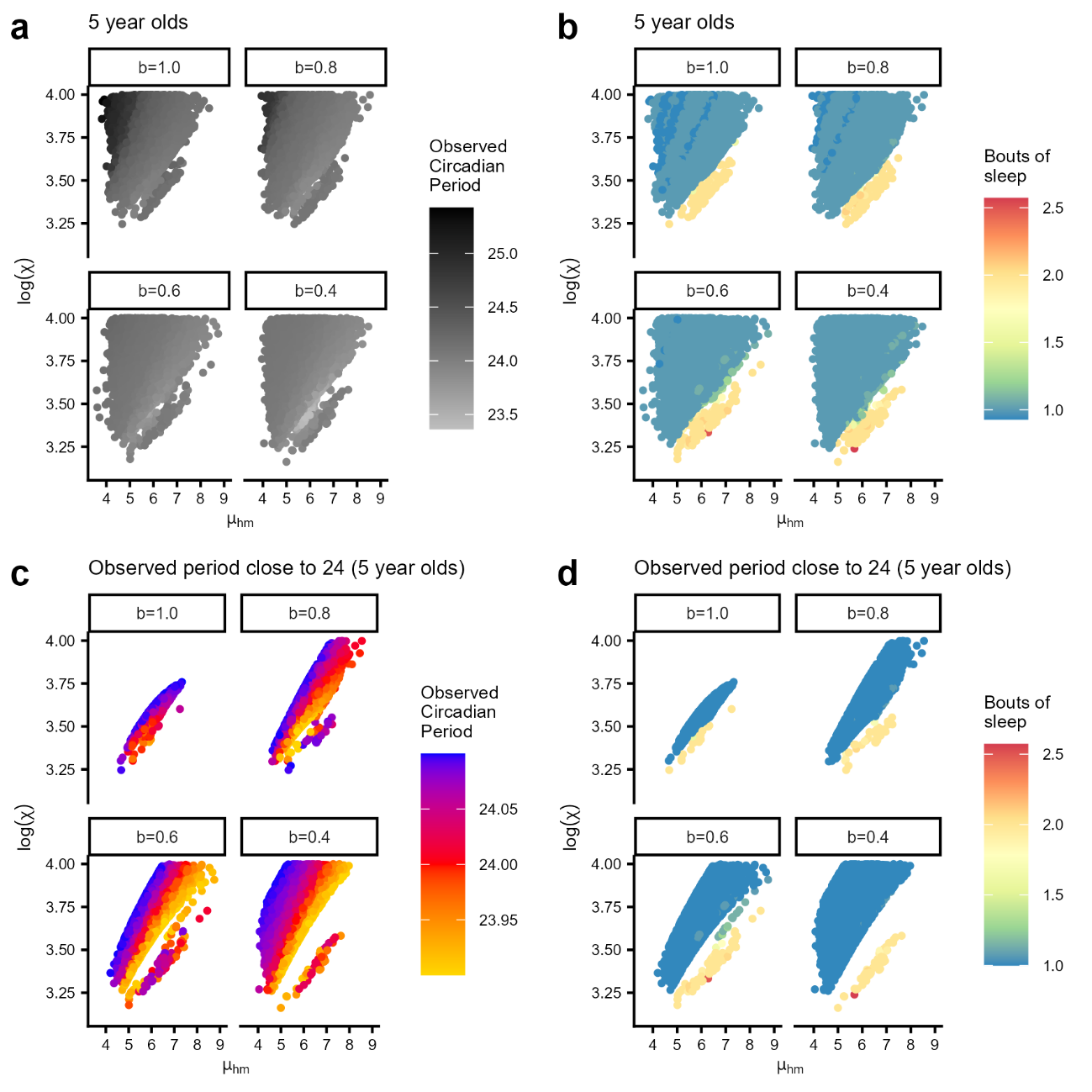

**Supplementary Figure S8**: The observed circadian period for in the $\mu_{hm}-\log\chi$ space in the 5-year-old case. **a)** The points from each posterior colored by the observed circadian period, showing the points with the highest period focused at low $\mu_{hm}$ and high $\log\chi$ in the b=1.0 case. The correspoding number of bouts per day in this parameter space is shown in **b)**, where divides between whole numbers of bouts (with a small amount of non-whole number between) is evident. Focusing on just the parameter combinations with period close to 24 h (<24.1 h and >23.9 h) gives small clusters of points in the space, colored by **c)** observed circadian period, and **d)** number of bouts of sleep per day.

**Appendix S5**

**Sensitivity analysis of the Mixed Gaussian for bout number**

To assess how the form of the mixed Gaussian used for the empirical distribution for average number of bouts affected the results, we made a second version of the $D_{0}-\chi$ model of Appendix S2 with the standard deviation of each Gaussian being one quarter that of the original, that is, $\sigma=\mu/40$. The mean of each Gaussian in the mixed Gaussian is the same as the original presented in Figure 2 and Table 1. The posterior densities (Supplementary Figure S9a and b) have a similar spread to those in Supplementary Figure S1, though are more clustered, particularly $\log\chi$. This was expected with the empirical distribution having more peaks (red density curves Supplementary Figure S9e). Similarly, the joint distribution and contours (Supplementary Figure S9c) have a similar spread to the original $D_{0}-\chi$ model results, though the clustering of $\log\chi$ is evident in the density contour lines. The sleep characteristics from the posteriors deviate slightly from the empirical distribution (Supplementary Figure S9d and e). For the 1-9 month old ages, the distribution of average total sleep per 24 hours is slightly right shifted and left skewed, and the proportion of 1-bout sleep is over-represented in the 1, 3, and 5 year ages.

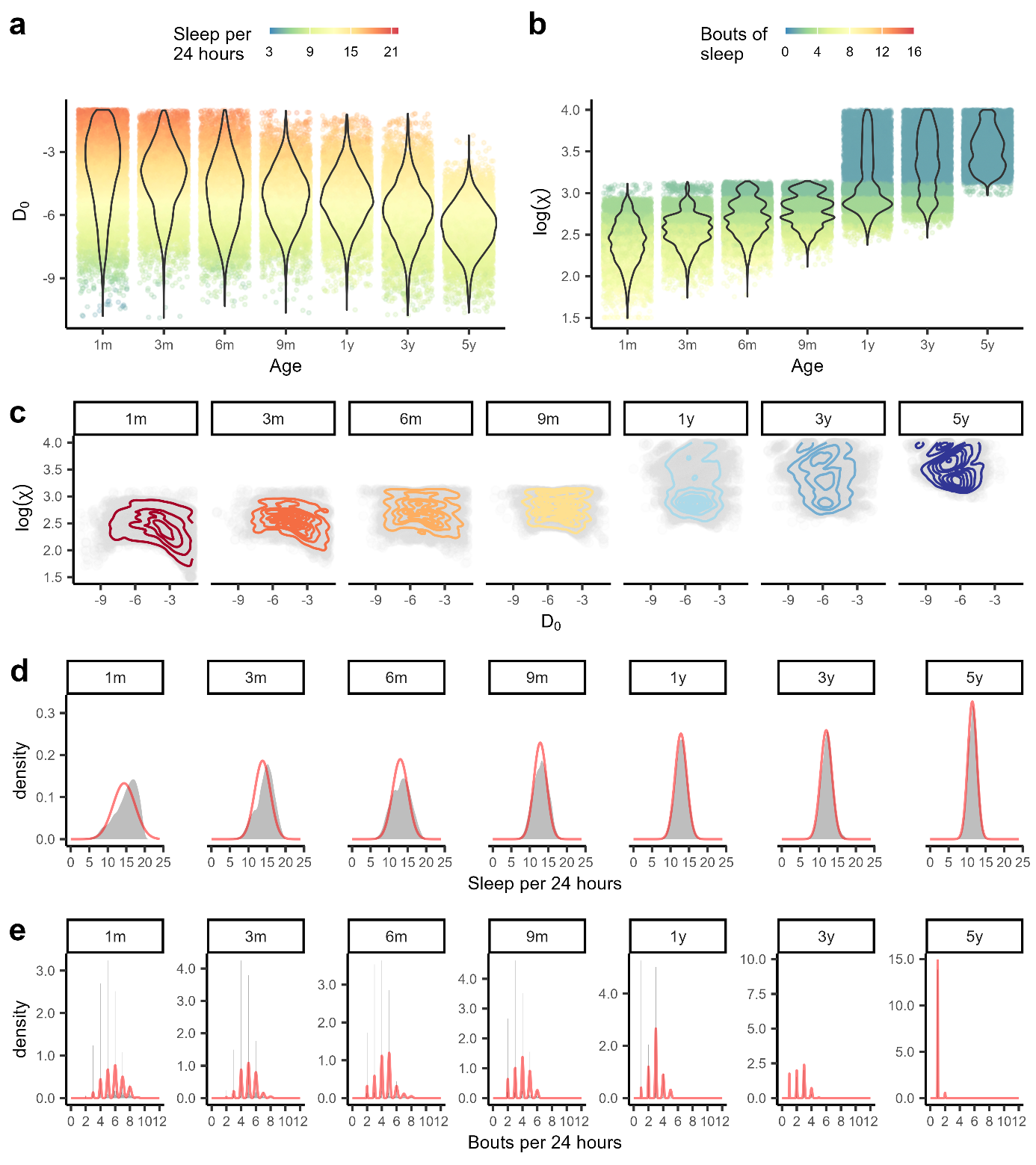

**Supplementary Figure S9**: Results from the $D_{0}-\chi$ model with different empirical distribution (Gaussians with smaller variances) for average number of bouts. Posterior distributions for each age for **a)** $D_{0}$ with corresponding average sleep per 24 hours, and **b)** $\log\chi$ with corresponding number of bouts of sleep. **c)** The joint posterior distribution for each age with density contours shown. The sleep characteristic distributions arising from the parameter posteriors, superimposed with the empirical density curves, are shown for **d)** amount of sleep per 24 hours and **e)** average bouts per 24 hours.
